# Extended-Ensemble Docking to Probe Evolution of Ligand Binding Sites During Large-Scale Structural Changes of Proteins

**DOI:** 10.1101/2021.03.28.437371

**Authors:** Karan Kapoor, Sundar Thangapandian, Emad Tajkhorshid

## Abstract

Proteins can sample a broad landscape as they undergo conformational transition between different functional states. As key players in almost all cellular processes, proteins are important drug targets. Considering the different conformational states of a protein is therefore central for a successful drug-design strategy. Here we introduce a novel docking protocol, termed as extended-ensemble docking, pertaining to proteins that undergo large-scale (global) conformational changes during their function. In its application to multidrug ABC-transporter P-glycoprotein (Pgp), extensive non-equilibrium molecular dynamics simulations employing system-specific collective variables capturing the alternate access mechanism of Pgp, are first used to construct the transition cycle of the transporter. An extended set of conformational states representing the full transition between the inward- and the outward-facing states of Pgp, is then used to seed high-throughput docking calculations of a set of known substrates, non-substrates, and modulators of the transporter. Large differences are observed in the predicted binding affinities to the conformational ensemble, with compounds showing stronger binding affinities to intermediate conformations compared to the starting crystal structure. Hierarchical clustering of the individual binding modes of the different compounds shows all ligands preferably bind to the large central cavity of the protein, formed at the apex of the transmembrane domain (TMD), whereas only small binding populations are observed in the previously described R and H sites present within the individual TMD leaflets. Based on the results, the central cavity is further divided into two major subsites: first subsite preferably binds smaller substrates and high-affinity inhibitors, whereas the second one shows preference for larger substrates and low-affinity modulators. These central sites along with the low-affinity interaction sites present within the individual TMD leaflets may respectively correspond to the proposed high- and low-affinity binding sites in Pgp. We propose further optimization strategy for developing more potent inhibitor of Pgp, based on increasing its specificity to the extended ensemble of the protein instead of using a single protein structure, as well as its selectivity for the high-affinity binding site. In contrast to earlier in-silico studies using single static structures of Pgp, our results show better agreement with experimental studies, pointing to the importance of incorporating the global conformational flexibility of proteins in future drug-discovery endeavors.

## Introduction

Modern drug discovery is centered around the identification of suitable protein targets that play important roles in a specific disease, and the development of small molecules designed to either attenuate or promote the function of those target proteins. ^1^ Structural determination techniques such as x-ray crystallography and cryo-EM have led to a large number of structurally known proteins that can help guide the drug discovery process. ^2, 3^ Experimentally resolved structures generally represent the starting or end states in the functional cycle of a protein, as the conformational intermediates (usually shorter-lived)arising during the function are often inaccessible under the experimental conditions. ^4^ This reduces the possible structural diversity of the target that can be utilized in designing more efficient drug discovery strategies. Generating and targeting these conformational intermediates still remains a major challenge in successful drug discovery campaigns.

Computational methods are increasingly used to complement the experimental drug discovery strategies. ^5, 6^ Structure-based drug discovery methods like docking utilize the available three-dimensional structure(s) of the target protein to make predictions for the best binding ligand candidates. ^7, 8^ The approach can significantly reduce the number of chemotypes to experimentally synthesize and test, compared to a high-throughput screening approach utilized alone without any prior knowledge of the potential interactions between the target and the small molecules. ^9^

Traditionally, and given the structural determination challenges described above, only a single experimental (e.g., crystal or cryo-EM) structure or a single computational model (e.g., a homology model) of the target protein has been used for docking purposes, in an approach we refer to as *s*ingle-point docking^10^ (Fig. 1). As stated above, these structures generally represent only a static snapshot of one functional state of the target protein. More recently, the single-point docking approach has been improved to take into account thermal fluctuations of the target protein in an approach referred to as *e*nsemble docking; the approach has been shown to further enhance both the quality and the efficiency of lead predictions ^11^ and successfully applied to a number of protein targets^12–15^ (Fig. 1). Ensemble docking makes use of molecular dynamics (MD) simulations for sampling the local conformational basin of the target in the vicinity of a starting experimental protein structure.^16–19^

**Figure 1:**
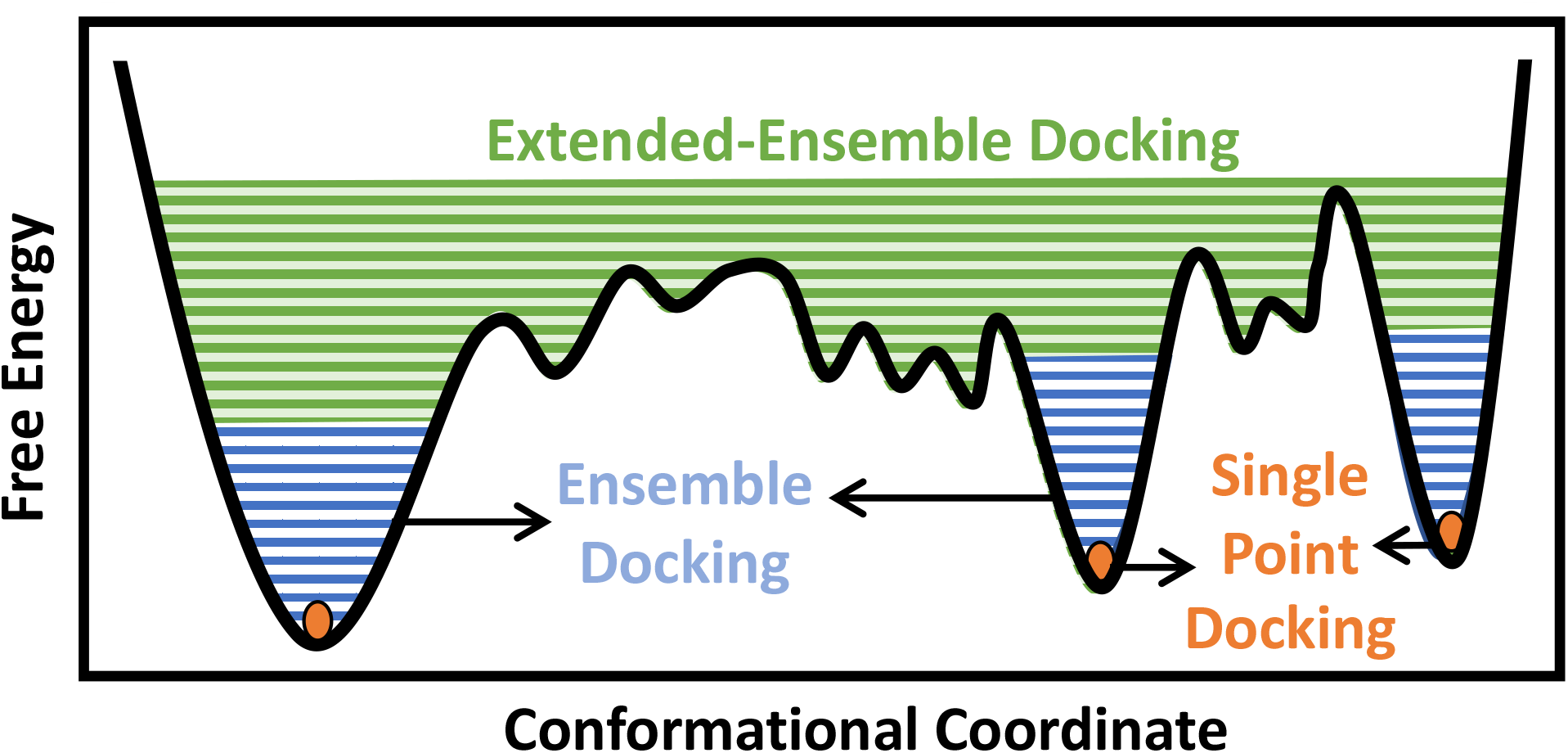
Conformational domains targeted by different docking approaches. Single-point docking utilizes a single structure of the protein target, restricting the sampling to a single point (orange dots) in the conformational landscape of the protein. Ensemble docking utilizes an ensemble of protein structures, often generated using MD simulations, taking into account thermal fluctuations within a local conformational basin in the vicinity of the starting experimental structure (blue lines). Extended-ensemble docking, the method introduced here, aims at taking into account the full functional cycle of the protein, generated, e.g., through the application of biased techniques to transition between the major functional states of the protein (green lines).

In the case of larger and more flexible protein targets though, which often display a complex conformational landscape with multiple local minima, ^4, 20^ the limited timescales of classical equilibrium MD simulations may not cover the large conformational heterogeneity of the target protein. ^21^ Large-scale motions and state transitions are of high relevance to the function of many macromolecular systems, e.g., the transmembrane transport of substrates by membrane transporters, activation of receptors, and gating of channels. ^4, 22^ Non-equilibrium methods allow us to expand our sampling of the phase space and to visit intermediate conformations formed during the transition between the major functional states of a protein. ^23, 24^ This forms the basis of what we term as *extended-ensemble docking*, that uses the enhanced sampling offered by non-equilibrium MD for generating an extended ensemble of the protein, sampling the full conformational transition pathway between its functional end states (Fig. 1). The extended ensemble of the protein can then be used to seed high-throughput docking calculations for specific drug-discovery applications.

Here we report the first application of this approach to investigation of ligand binding in the ABC exporter, P-glycoprotein (Pgp),^25^ a transporter that is over-expressed in the plasma membrane of cancer cells and a major cause for the development of multi-drug resistance. ^26, 27^ The transporter accomplishes its function as a cellular ‘vacuum cleaner’ by binding different classes of molecules ^28, 29^ through a large central drug binding pocket (DBP), formed by its two opposing, pseudosymmetric transmembrane domains (TMDs) (Fig. S1). As an active transporter, Pgp has to follow the general ‘alternating-access’ model ^30, 31^ for its function, thus alternating between the inward-facing (IF) and outward-facing (OF) states, alternatively exposing the DBP (and the bound substrate) to the intracellular and extracellular sides of the membrane. The large-scale structural transitions of the transporter are fueled by and coupled to ATP-driven dimerization and opening of its two nucleotide binding domains (NBDs) connected to the TMDs^32–34^ (Fig. S1).

A number of computational studies employing molecular docking, ^35–37^ pharmacophore mapping, ^38^ machine learning, ^39^ and MD-based approaches ^40^ have been conducted to characterize better the promiscuity of the binding sites in Pgp. However, these studies have generated vastly conflicting results to each other and to earlier mutational studies; ^41–43^ it still remains debatable as to whether diverse molecules bind to distinct or overlapping binding sites in Pgp. A major limitation of the in-silico approaches for the case of Pgp is that they neglect the structural heterogeneity inherent to a flexible, multidomain protein such as Pgp by targeting only a single crystal structure/homology model, providing only a snapshot and ignoring the myriad of conformations arising during the function of the protein.

In the present study, we use non-equilibrium, driven MD simulations employing system-specific reaction coordinates describing the alternate-access transitions of ABC transporters, in order to sample the full transition cycle of Pgp between the IF and OF states. Docking of a set of small molecules, including substrates, non-substrates, and modulators, to the generated *e*xtended ensemble of the protein reveals that different classes of compounds may bind distinctly to different conformational states/intermediates of Pgp. The captured differential ligand-binding properties of different “subsites” in Pgp, whose accessibility and size are modulated by the protein conformational changes as the protein undergoes transition between the IF and OF states, may be a determining factor in the broad substrate specificity of the transporter. Our results show a close relationship with the previous experimental studies and additionally allow us to make suggestions for developing more potent inhibitors of this multidrug resistance transporter.

## Computational Approach

The main distinction of extended-ensemble docking from other docking approaches is to first generate structural models for the intermediate conformational states of the protein that arise during the transition between its structurally known states, e.g., through the application of biased simulation techniques. Then, instead of targeting a single experimental/modeled structure, the entire protein extended ensemble is targeted through high-throughput docking calculations (extended-ensemble docking). Following are the details of the different steps involved in the protocol used to generate the target conformational ensemble and its specific application to Pgp (Fig. 2). We also present a suitable method for clustering the predicted binding modes of compounds to the ensemble of structures, in order to obtain physiologically relevant results that are related to the mechanism of the protein.

**Figure 2:**
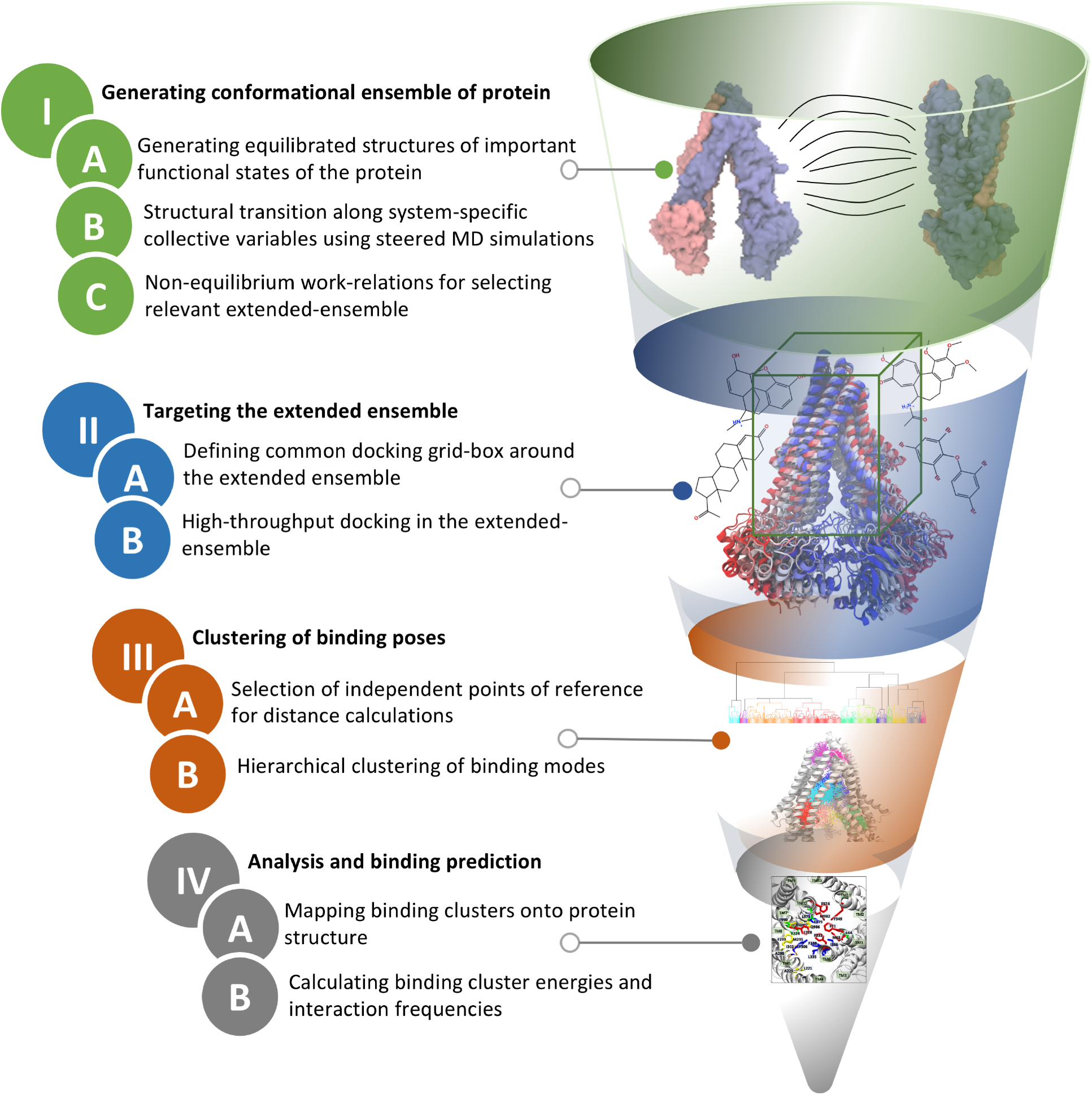
Extended-ensemble docking protocol. Flow diagram showing different steps involved in the *e*xtended-ensemble docking approach. The approach involves the targeting of an extended ensemble of the protein, generated along its functional cycle, by docking small molecules, followed by clustering of the predicted binding poses for each representative conformation. See Methods for details of each step.

### I. General strategy to generate an extended conformational ensemble for the target protein

We start with the available (known/modeled) structures of major functional states (end states) of the protein. In order to obtain relaxed structures, which are required for optimal sampling of the transition pathway between them, one would need to first equilibrate them in a native environment, e.g., solution for globular proteins or a lipid bilayer for membrane proteins.

Subsequently, in order to describe the conformational transition between major states in a physiological context, we define a set of system-specific collective variables (CVs) that capture the essential conformational changes in the protein during the transition. These CVs need to be closely related to important/relevant structural features of the different states of the protein; by applying time-dependent biases along these CVs, for example by using biased methods such as steered MD (SMD),^44^ we can generate transition pathways connecting the end states. ^45, 46^ The amount of non-equilibrium work needed to induce the conformational change between two states during a driven transition along a particular pathway (obtained from multiple simulation replicates) provides an approximate initial metric for the likelihood and relevance of the sampled transition pathway.^47, 48^ At the same time, the RMSD of the resulting structure from the target structure provides a measure for the effectiveness of the prescribed protocol to complete the transition. Low RMSD to the target structure and low non-equilibrium work values for a pathway point towards the efficiency and the quality of the applied transition pathway/protocol. The conformational ensemble of the protein can then be generated by selecting structures spanning the distance (along the CV space) between the two end states, for example, by uniformly selecting representative conformations along the optimal transition pathway.

In the following sections we describe each of these steps for the specific example of Pgp.

#### Equilibration of functional states of Pgp

In the case of Pgp, crystallographic studies have captured only the IF state of the transporter thus far. ^28, 49–52^ Additionally, these crystal structures show different degrees of NBD separation, pointing to the conformational heterogeneity of the transporter in the IF state. Conventional MD simulations were thus employed first to allow further sampling of the (local) conformational space of the IF state. We had previously generated an equilibrated ensemble of Pgp IF structures, ^53^ using multiple, independently run MD simulations of the crystal structure of mouse Pgp (PDB: 4M1M^28^) fully embedded in POPC/cholesterol lipid bilayers. As binding of ATP/Mg^2+^ to the NBDs of the transporter is a prerequisite for their dimerization and progression of the catalytic cycle, ATP/Mg^2+^ were carefully modeled into their respective binding sites in the NBDs following the protocol described by Wen et al, ^54^ that reproduces the conserved nucleotide-binding characteristics observed in high-resolution ABC transporter structures.

As for the OF state, recently a cryo-EM structure of Pgp in this state was published. ^55^ Due to a Q/E mutation at the catalytic site of the structure, it has been suggested to be a ’dead-mutant’, a non-catalytic form of Pgp incapable of substrate transport. ^56^ Other structural studies have instead suggested this structure to represent a post-transport, collapsed-occluded state of the transporter. ^57^ Additionally, studies using DEER spectroscopy have shown that Pgp samples a wider opening on the extracellular side of the TMDs in its OF state during the catalytic cycle, ^58^ not seen in the cryo-EM structure. Due to these reasons, we have generated our own equilibrated model of Pgp OF state using careful sequence alignment, homology modeling, and SMD simulations, and tested its stability in multiple MD simulations. ^58^ The distances between different regions of the transporter present in this state were found to match well with the spectroscopy studies ^58^ described above.

#### System-specific CVs for sampling conformational transition

In our previous work with ABC transporters, including Pgp, we developed a set of specific CVs to capture the motion associated with the alternate-access mechanism of these transporters. ^48, 58^ Conformational transition between the equilibrated IF and OF states of Pgp was carried out using these system-specific CVs, controlling both the global conformational changes of the protein during the transition, as well as local interactions (e.g., protein-protein interactions at the interface) crucial for the formation of stable Pgp states (Fig. S2). These include two orientation-based CVs (quaternions), denoted as *α* and *β*, describing the closing/opening of the TMDs on the intracellular and extracellular sides, respectively, during the IF-OF transition. These specific orientation quaternions are defined on the basis of the C*α* positions of the two TMD bundles in IF and OF structures, respectively, and are associated with the relative rotation of the helices during the transition between the two states (see Verhalen et al^58^ for more details). Additionally, one CV, denoted as SB, controls the formation of a salt-bridge interaction (K185-D993) in the middle of the TM helical region that may stabilize the cytoplasmic closure of the TMD in the OF state. Furthermore, for proper dimerization of the NBDs during the formation of the OF state, 12 CVs (collectively denoted as NBDi) were used to enforce distances between different atoms of ATP in each NBD and the signature motif (LSGGQ) of the opposing NBD, and between NBDs X-loops and coupling helices located at the interface between the TMD and NBDs (Table S1). All the target distances for NBDi CVs were obtained from the high-resolution crystal structure of the dimerized NBDs of HlyB (PDB: 1XEF^59^), an ABC exporter.

#### Driving structural transition using SMD simulations

We performed SMD simulations employing different protocols, each defined by applying a distinct order of the four system-specific CVs described above (*α*, *β*, SB and NBDi), to explore a wide-range of mechanistically distinct transition pathways in Pgp. The transition protocols included: P1: *α* + NBDi + SB *→ β*, P2: NBDi *→ α* + SB *→ β*, P3: *α* + SB *→* NBDi *→ β*, and P4: *α* + NBDi + SB + *β*, where ‘+’ indicates that the CVs are applied simultaneously, and ‘*→*’ denotes their sequential application.

SMD simulations were performed starting from 20 equilibrated ATP/Mg^2+^-bound IF structures of Pgp that were already embedded in POPC/cholesterol lipid bilayers, solvated and neutralized with Na^+^ and Cl*^−^* ions, ^53^ and targeting the equilibrated model of the OF state, ^58^ using the 4 transition protocols (P1, P2, P3 an P4). This amounted to a total of 80 (20 starting structures *×* 4 transition protocols) independent SMD runs. All simulations were performed using NAMD2^60, 61^ with the CHARMM36 force-field representing protein, lipid, and nucleic acids^62–64^ and TIP3P model for water. ^65^ The simulations were carried out in an NPT ensemble, with the temperature maintained at 310 K using Langevin dynamics^66^ with a damping coefficient of *γ* = 0.5 ps*^−^*^1^, and the pressure maintained constant at 1 bar using the Nośe-Hoover Langevin piston method. ^67, 68^ Periodic boundary conditions were used, and Long-range electrostatic forces were calculated using the particle mesh Ewald (PME) method. ^69^ The non-bonded interactions were calculated with switching and cutoff distances of 10 Å and 12 Å, respectively. A 2 fs integration timestep was used for calculating the forces. Eighty independent SMD runs were performed for 30 ns each to bring the starting structure close to the target orientations and distances (Table S1). Harmonic force constants of 10^5^ kcal/mol/rad^2^ and 2 kcal/mol/Å^2^ were used for the quaternion-based and distance-based CVs, respectively. Subsequently, non-equilibrium work profiles and C*α* RMSD with respect to the target OF structure were calculated using our in-house TCL scripts in VMD.^70^

**Table 1:**
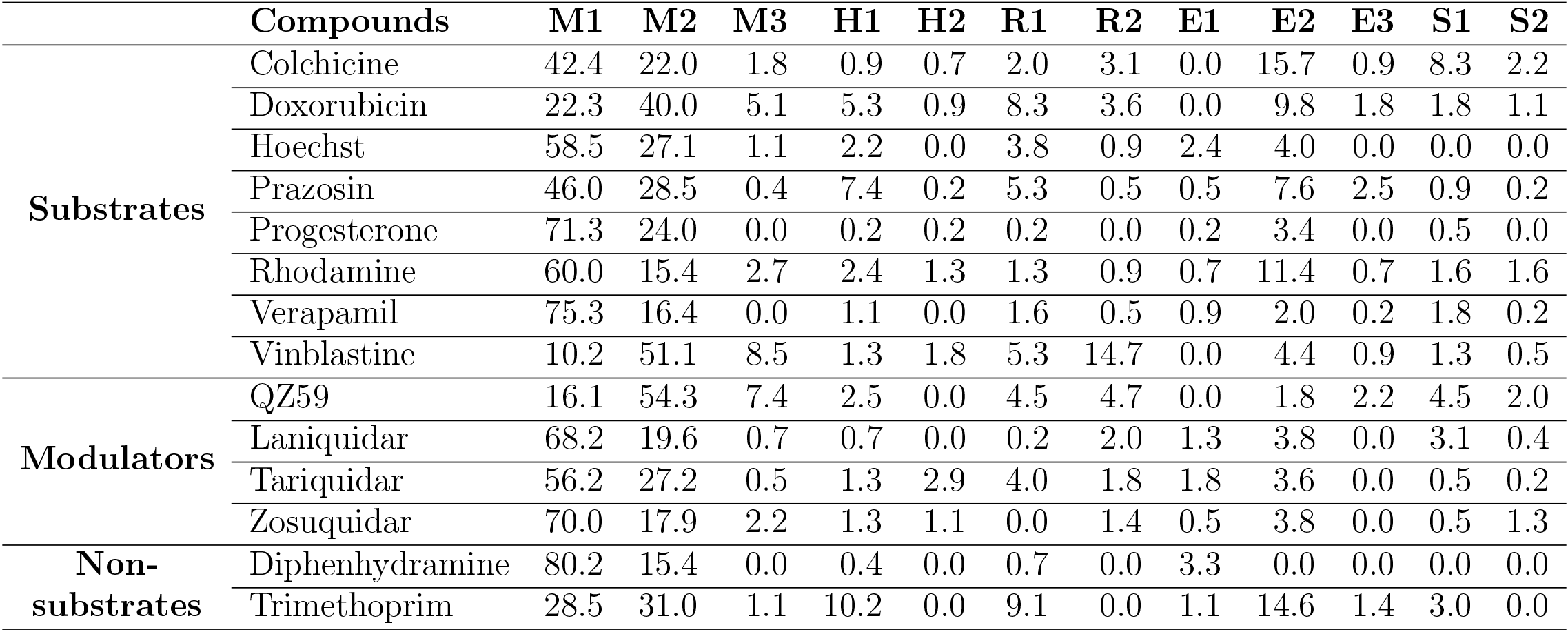
Distribution (percentage) of binding modes of different compounds to different binding sites observed in the extended ensemble of Pgp.

#### Using the lowest-work transition pathway to select an extended ensemble

In the next step, starting with an equilibrated IF structure with largest RMSD to the target OF structure we run a longer (100 ns) SMD simulation using the most efficient transition protocol (lowest work value protocol from the set of shorter SMD runs described above). The trajectory of this run will be used to generate the extended ensemble of the protein. These longer (slower) SMD simulation decreases the non-equilibrium work required for the transition and allows the protein to remain closer to a low-energy path during the transition. An ensemble of the protein conformations was then extracted by taking 50 snapshots from the run distributed nearly equally along the CV phase-space (equally spaced in time). We also analyzed the conformational modulation of the putative binding pockets in the Pgp ensemble over the course of the transition using `Site Finder` in Molecular Operating Environment (MOE),^71^ that calculates both the size and the ligand binding propensity (based on the amino acid composition) of the predicted binding pockets (also termed active binding pockets).

### II. Targeting the extended ensemble by molecular docking

In extended-ensemble docking, we aim at utilizing the conformations and intermediate states generated along the global transition pathway of a protein for docking. The selection of a suitable docking methodology generally depends on the specific goal of the study. In the following section we describe the steps involved in the docking calculations carried out in the extended ensemble of Pgp.

In order to compare the results obtained from our extended-ensemble approach with other in-silico studies, we used a previous single-point docking study^36^ conducted on the crystal structure of the IF state of Pgp (PDB: 4M1M^28^) as a point of reference. We obtained a set of 14 compounds for our docking study from this single-point study that comprised different classes of compounds (substrates, non-substrates and high- and low-affinity modulators) known to bind (and in the case of non-substrates, not bind) to Pgp. Docking of compounds, while allowing flexibility around all rotatable bonds, was carried out using Autodock Vina^72^ in the extended ensemble of pgp along with the starting crystal structure of the IF state (Fig. 3). All structures were superimposed on the starting crystal structure, centered at (0, 0, 0), to discount for the translation and rotation of the protein during the simulation. A common docking grid-box was then defined around the extended ensemble covering the TMD apex (putative binding sites) of the protein. This box centered at (0, 0, 40) had dimensions of 80, 54 and 80 Å in the *x*, *y* and *z* directions, respectively (Fig. 3). Docking of compounds was carried out in this grid-box in an agnostic manner, i.e., the compounds were allowed to sample the entire binding space within the box without any constraints. A total of nine binding poses were generated for each docked compound in each protein conformation.

**Figure 3:**
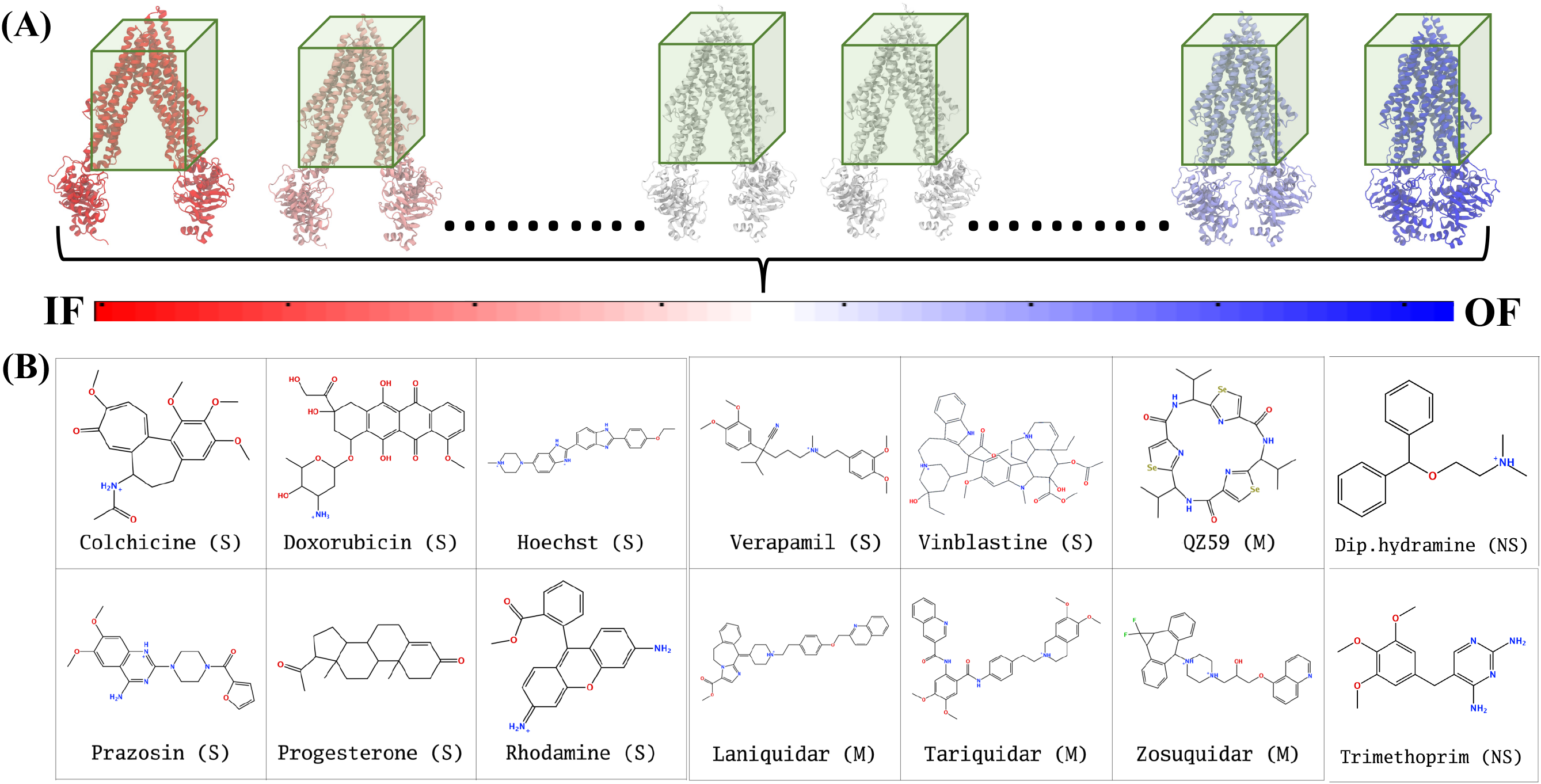
Docking in extended ensemble. (A) The extended ensemble of Pgp (with color changing from red to blue between states depending on the position along the trajectory), generated by taking 50 snapshots nearly equally distributed along the CV phase-space defining the IF to OF transition. Molecular docking was carried out in a docking grid-box (shown in green) defined around the TMD of the protein in all conformations. (B) The chemical structures of the 14 compounds selected for docking in the extended ensemble of Pgp are shown. These compounds include known substrates (S), modulators (M) and non-substrates (NS) of Pgp.

### III. Clustering of binding poses

Clustering of the predicted binding poses can be challenging due to the large-scale structural differences of the protein conformations in the extended ensemble. For example, clustering of the docked compounds based on simple distance-based metrics, e.g., RMSD between the docked compounds, can misleadingly generate separate binding clusters for compounds that bind to the very same region of the protein. To overcome this problem, we first calculate the distances of each docked compound with respect to a hybrid system of reference, combining protein-independent and -dependent frames of reference, defined in the three-dimensional space and then used these distances as clustering inputs. The clustering strategy utilized here can be generalized to other cases where one deals with similarly flexible protein targets.

Distances of the predicted binding modes of each docked compound in the extended ensemble of Pgp were calculated with respect to 6 reference points in the three-dimensional space: two variable points in the +*x*, -*x* directions and four fixed points in the +*y*, -*y*, +*z*, -*z* directions, respectively. As the closure of the intracellular end of the TMDs in the *±x* direction drives the conformational changes in the IF to OF transition, selection of variable *±x* points located on the protein helices involved in the conformational change, and fixed points in the *±y*/*±z* directions, sufficiently allows the generation of similar distance values for the compounds binding to the same regions of the protein (Fig. S3). We used the same docking grid-box defined around the TMD region (centered at (0, 0, 40) with dimensions of 80, 54 and 80 Å in the *x*, *y* and *z* directions, respectively), four fixed points in the *±y*/*±z* directions were selected as the centers of the four side faces of the box, lying at (0, 27, 40), (0, -27, 40), (0, 0, 0) and (0, 0, 80), respectively. The two variable points in the *±x* directions were selected as the C*α* atoms of the first residues of the TM1 (L45) and TM7 (P705), respectively. Subsequently, the distances of the center of mass of the docked compound from these reference points were used for clustering.

The above six distances used to define the position of each docked compound were used for clustering the docked poses by hierarchical clustering method in MATLAB.^73^ In the first step, a dissimilarity matrix between each pair of objects (distances) in the data set was calculated. The data set for all compounds, consisting of 6,300 distances (50 protein conformations *×* 14 compounds *×* 9 binding modes for each compound) thus generated a dissimilarity matrix consisting of (6, 300 *×* (6, 300 *−* 1))*/*2 pairs. In the second step, a hierarchical tree was constructed by linking pairs of objects in close proximity (based on the dissimilarity matrix) using the average linkage method. ^74^ In the third step, a distance cutoff was used for pruning the branches of the hierarchical tree so obtained, allowing partitioning of all points lying below a specific branch into separate clusters. Using trial and error, a distance cutoff of 15 was selected, providing a suitable partitioning of the hierarchical tree, with the different pruned branches, representing the different clusters, found to bind to different regions of the protein.

### IV. Analysis and binding predictions

The binding of different compounds to the extended ensemble of the Pgp was characterized in terms of the predicted binding affinities for the different binding modes of each compound in each cluster, as well as the relative population distribution of these compounds in each cluster. Mapping of these clusters onto the TMD of the protein allowed identification of the corresponding binding sites in the protein along with the residues displaying the highest overall contact frequencies with each compound. In the latter case, any protein heavy atom within 4 Å of a heavy atom of the ligand was considered to form contacts.

## Results

Here we describe the results for the application of the extended-ensemble docking approach to the multi-drug transporter Pgp. The results of generating the extended ensemble of Pgp using SMD simulations are provided in Supplementary Information (Fig. S4, S5, S6, S7 and S8). Here, we first describe the results evaluating the conformational modulation of the possible binding sites formed in Pgp as it transitions from the IF to OF state. Subsequent docking calculations carried out to the selected ensemble of the protein allowed us to further characterize differential binding affinities displayed by compounds in the binding sites present in the different conformations of Pgp. Furthermore, these calculations allowed us to differentiate between the different classes of compounds based on their binding preference for specific sites in the DBP of Pgp.

### Evolution of binding pockets in Pgp during IF to OF transition

The analysis of the physicochemical properties of the binding pockets appearing in the extended ensemble of Pgp, as it transitions from the IF to the OF state, allowed us to evaluate the changes in these binding pockets due to the conformational changes in the protein. Three binding pockets are formed in the membrane encompassing region of the TMDs that comprises the central DBP in Pgp (Fig. 4). Out of these, the pocket at the TMDs apex (where the two TMD leaflets meet; shown in blue in Fig. 4) displays a high ligand binding propensity in all conformations except in snapshot 50 representing an OF state (Table S2). This observation can be expected as this pocket formed in the OF state undergoes significant shape change due to rearrangement of TMD helices (Fig. 4G). The largest size for this pocket (along with the highest ligand-binding propensity) is observed in snapshot 30 (Fig. 4E) which is an IF-like state conformationally similar (RMSD of 3.2 Å) to the starting crystal structure (Fig. 4A). The other two binding pockets formed in the DBP lie within the individual TMD leaflets (shown in red and green in Fig. 4), and show relatively smaller sizes and lower ligand-binding propensities in different protein conformations compared to the apex binding pocket. Additional small binding pockets exhibiting low ligand-binding propensities are also observed to form in the late conformations of Pgp extended ensemble, either on the intracellular side, where the two TMD leaflets come together and close the cytoplasmic end of the protein lumen (shown in pink in Fig. 4F) or on the extracellular side, where the two TMD leaflets separate and lead to extracellular opening (shown in yellow in Fig. 4F and G).

**Figure 4:**
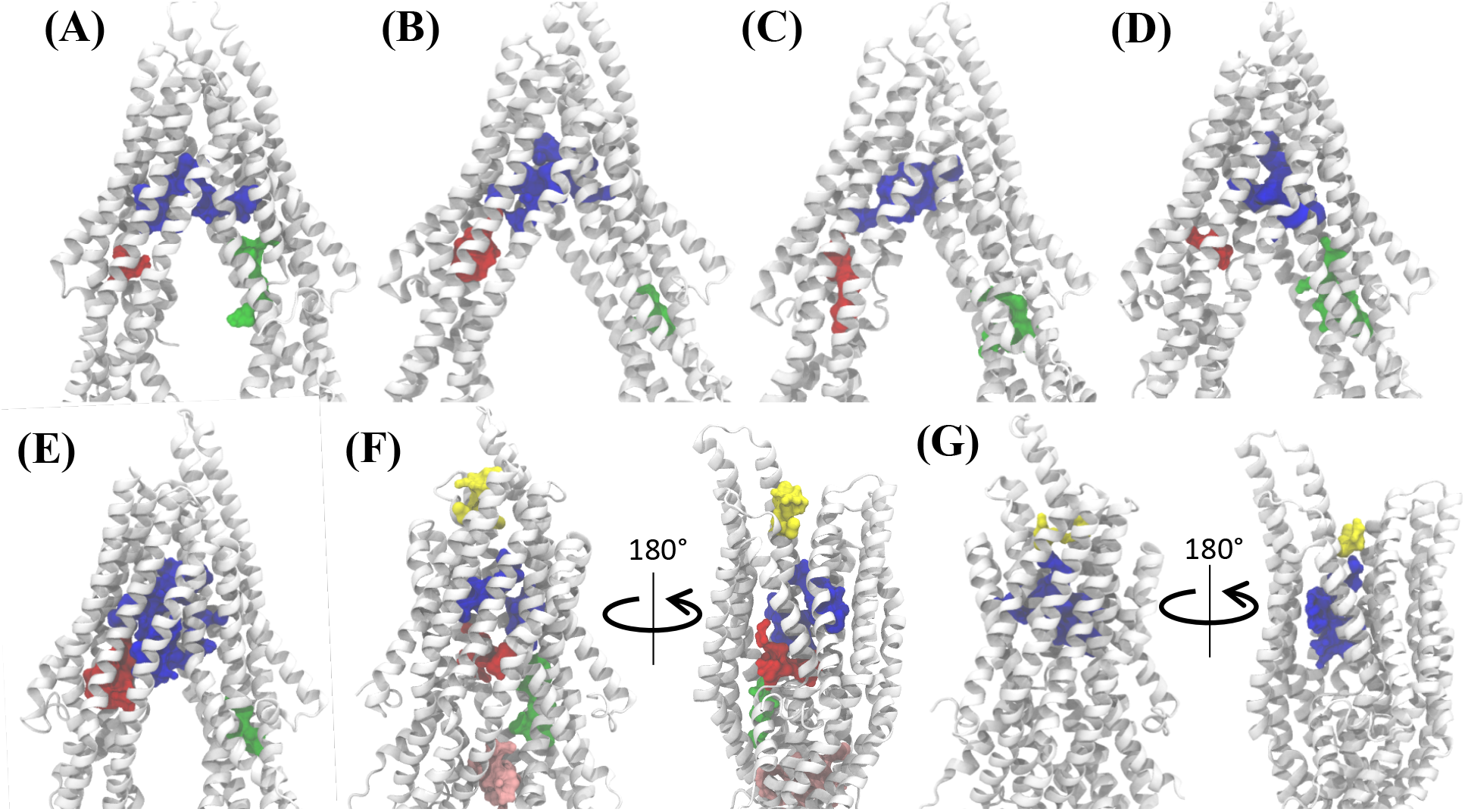
Binding pocket predictions during Pgp IF to OF transition. A) The active binding pockets predicted in the starting IF crystal structure are shown. The different binding pocket residues are shown by space filling representations in different colors. B-G) Active binding pockets predicted in snapshots 1, 10, 20, 30, 40 and 50 of the extended ensemble of Pgp, respectively. The large central pocket (in blue) in the apex of the DBP shows the highest ligand binding propensity and is present in all protein conformations.

### Differential binding affinities of compounds predicted in different protein conformations

Docking of different compounds was carried out to an extended ensemble covering the full IF-to-OF transition of Pgp. The tested compounds were found to dock with different binding affinities to the different protein conformations (Fig. 5 and Fig. S9). Interestingly, all compounds show stronger binding affinities (more negative docking scores) in conformations different from the starting IF crystal structure, reaching binding affinities as much as -4.1 kcal/mol higher than the crystal structure (laniquidar shown in Fig. S9). In general, high-affinity Pgp modulators (laniquidar, tariquidar, zosuquidar) show stronger binding affinities than low-affinity modulators (QZ59), large substrates (doxorubicin, vinblastine) and small substrates (hoechst, verapamil, progesterone, prazosin, rhodamine, colchicine). Consistent with their known properties, non-substrate ligands (diphenhydramine, trimethoprim) show the weakest binding affinities among all compounds.

**Figure 5:**
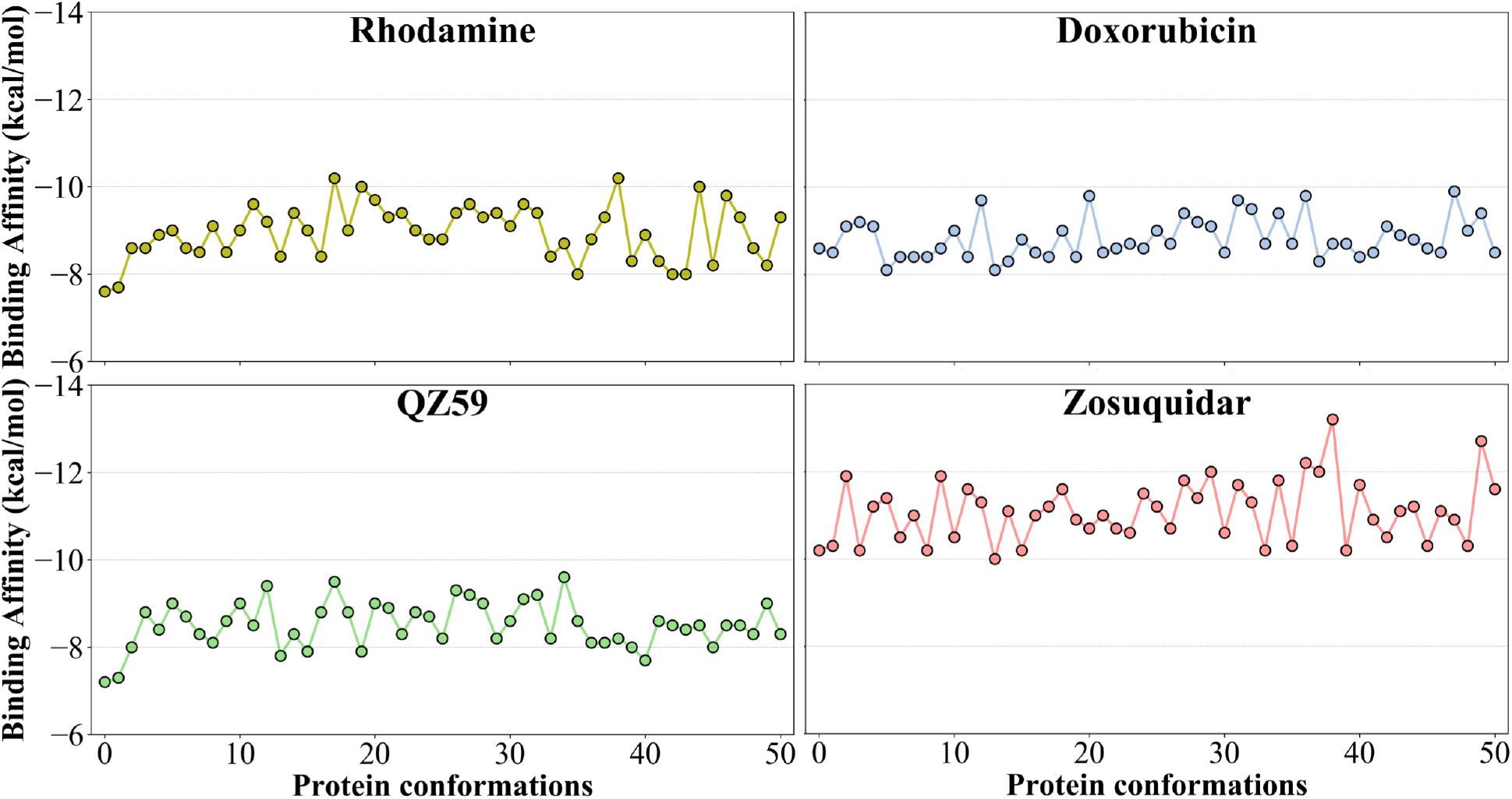
Binding affinities to the extended ensemble. The highest predicted binding affinities for 4 representative compounds (small substrate: rhodamine; large substrate: doxorubicin; low-affinity modulator: QZ59; high-affinity modulator: zosuquidar) to each conformation of the extended ensemble of Pgp are shown in different panels. Data for the rest of the compounds are shown in Fig. S9. Conformation 0 represents the starting IF crystal structure used in simulating the transition cycle, and the selected conformations along the IF to OF transition pathway are numbered 1 to 50. The predicted binding affinities fluctuate between different protein conformations, with the high-affinity modulator showing the highest binding affinities among all compounds.

### Preferential binding to two major sites

The hierarchical clustering of binding modes of the tested compounds, allowed us to characterize their preference for different binding regions of Pgp. Using a cutoff distance of 15 Å the resulting hierarchical tree was partitioned into 12 separate clusters (Fig. S10). The relative populations of the clusters showed that Clusters 4 and 5 together constituted *∼*78% of all predicted binding modes. In contrast, relatively small number of binding mode populations (2 *−* 7%) were observed in Clusters 3, 6, 7, 8, 10 and 11 and only marginal numbers of binding modes (*<* 1%) were observed in the other 4 Clusters (Table S3). Mapping of the different clusters onto the protein TMDs showed that Clusters 1, 2 and 3 (named E1, E3 and E2 sites, respectively) bind to the extracellular side of the TMDs, Clusters 4, 5 and 11 (named M1, M2, and M3 sites, respectively) bind to the apex of the TMDs, Clusters 6 and 7 (named R2 and R1, respectively) bind to TMD2, Cluster 8 (named S1) binds to the membrane-facing surface of TMD2, Cluster 9 (named S2) binds to the intracellular end of the TMDs, and Clusters 10 and 12 (named H1 and H2, respectively) bind to TMD1. The naming convention for binding sites assigned is further described in Discussion.

High-affinity modulators (laniquidar, tariquidar, zosuquidar) and small substrates (hoechst, verapamil, progesterone, prazosin, rhodamine, colchicine) show high binding populations to the M1 site, located at the apex of the TMDs, whereas the low-affinity modulator (QZ59) and large substrates (doxorubicin, vinblastine) show preferred binding to M2, which is still located at the apex but partially towards TMD2 (Fig. 6, S11 and Table 1). Overall, these two binding sites in the top half of the DBP show relatively stronger binding affinities also for other tested compounds The M3 site, below M1 and towards TMD1, is only sparsely populated for all compounds except large substrates and low-affinity modulators that show moderate populations in this site (Fig. 6, S11 and Table 1). This binding site is seen to form only during the late transition to the OF state of Pgp (Fig. 7 and S12). R1 and R2 sites are below the M2 site in TMD2, whereas H1 and H2 sites are present below the M3 site in TMD1 (Fig. 6 and S11). Low-affinity modulators and large substrates display moderate binding populations to the R1/R2 sites, while small substrates like prazosin show moderate binding to H1 (Table 1). The H2 site, on the other hand, shows marginal binding populations for all compounds. In general, R1 and H1 are seen to form in protein snapshots 1-30, while the protein is still in open IF-like conformations, whereas R2 and H2 sites are sampled more during the later phase of the transition (snapshots 30-50), with the protein present in OF-like conformations (Fig. 7 and S12). Extracellular sites E1 and E2 are found in OF-like conformations (snapshots 30-50), where the DBP opens up towards the extracellular side (Fig. 7 and S12). Most compounds are seen to bind with substantial populations to the E2 site, whereas E1 shows marginal binding (Fig. 6, S11 and Table 1). Some of the compounds also show binding to an additional extracellular site, E3, at the top and on the outer surface of the TMDs facing the extracellular environment. This site is present during the early phase of the protein transition (snapshots 1-30), i.e., in IF-like states, and hence it may not represent a physiologically important site for the substrates as the DBP is inaccessible from the extracellular side in these conformations.

**Figure 6:**
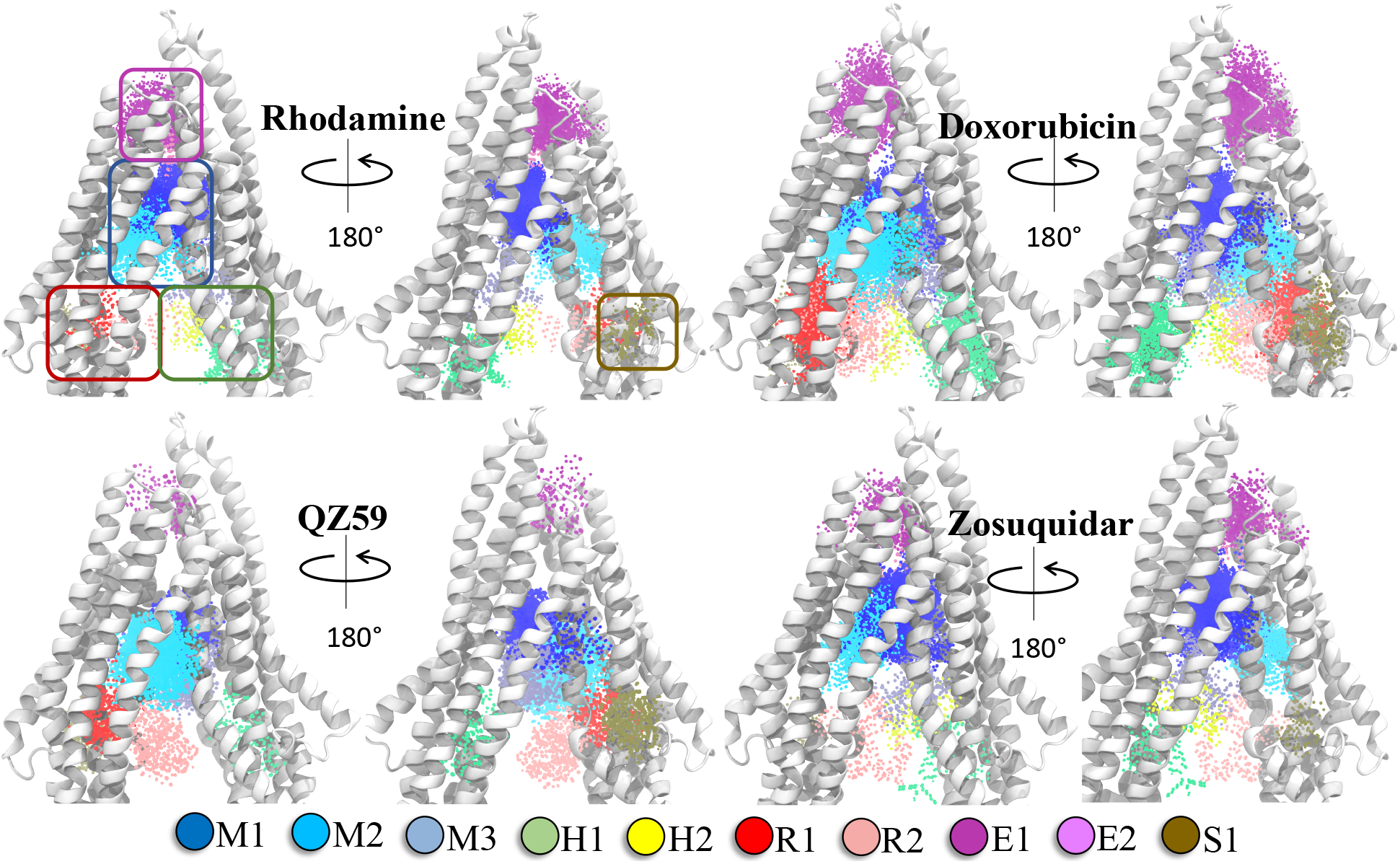
Clustering of binding modes generated from docking calculations. Clustering of the binding modes for 4 representative compounds to the extended ensemble of Pgp is shown (rest of the compounds are shown in Fig. S11). Only one representative, IF-like conformation is shown here for clarity. The main binding sites in the TMDs are marked by colored rectangles (blue: modulator or M site; green: hoechst-binding or H site; red: rhodamine-binding or R site; purple: extracellular or E site; brown: subsidiary or S site) shown for the first representative compound (rhodamine). Binding clusters (or binding subsites) within the main binding regions are highlighted with colored points (indicated in the legend at the bottom), representing the heavy atoms of the clustered binding modes (E3 and S2 sites are not shown as they may not represent sites important for substrate binding/transport). The density of points in each cluster represents the cluster population. M1 and M2 subsites present at the apex of the TMDs show the highest cluster populations for all compounds.

**Figure 7:**
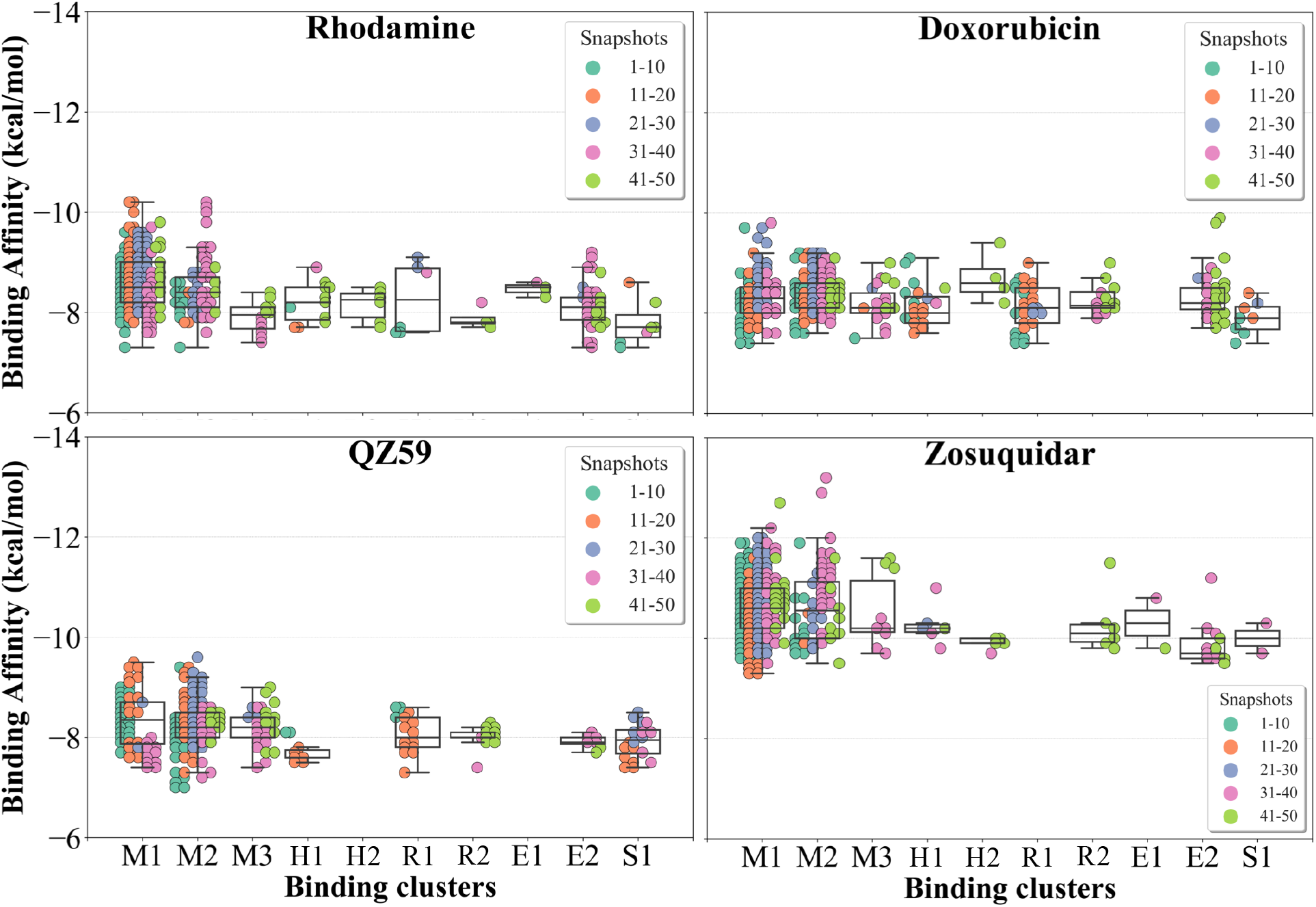
Binding cluster energies. The binding affinities of members (binding modes) of each binding cluster (binding subsites) are shown as a swarm plot for 4 representative compounds (data for the rest of the compounds are shown in Fig. S12). The snapshots (1-50) representing protein conformations as it transition from the IF to OF state shown in different colors (defined in legend). Additionally, a boxplot providing the median cluster values, Q1 and Q3 quartiles, as well as minimum (Q1 *−* 1.5 × interquartile range) and maximum (Q3 + 1.5 × interquartile range) binding affinity values, is overlaid on top of the swarm plot for each cluster. M1 and M2 binding clusters show the highest binding populations and display the strongest binding affinities for all compounds.

Other than the above ten binding sites, some compounds show binding to two additional subsidiary sites S1 and S2, with S1 on the membrane-facing surface of TMD2 and formed between the elbow helix and the broken (unstructured) region of TM12 (Fig. 6, S11 and Table 1), and S2 at the bottom of the TMDs between TM4 and TM6. S2 is seen to form only in the later protein conformations (snapshots 40-50), i.e., in OF-like states, and, similar to E3, may not represent a physiologically important site as the DBP may become inaccessible in these conformation due to the closing of the TMDs on the intracellular side during the IF-OF transition.

### Important binding residues in major sites

The contacts formed by the protein residues with each compound were further analyzed in terms of interaction frequencies. Comparatively, high-affinity modulators (laniquidar, tariquidar and zosuquidar) and some substrates (progesterone, verapamil) show higher interaction frequencies with the same set of residues than low-affinity modulator (QZ59) and both large (doxorubicin, vinblastine) and small substrates (colchicine, hoechst, prazosin, rhodamine) (Fig. 8 and Fig. S13). The non-substrate trimethoprim shows the lowest interaction frequency among all compounds, relating to its low selectivity for Pgp. Overall, most com-pounds form highly favorable interactions with residues in TM1, TM5, TM6, TM7, TM11 and TM12 that come together to form the DBP, with the exception of large substrates and the low-affinity modulator that instead display higher interactions with TM4 residues.

**Figure 8:**
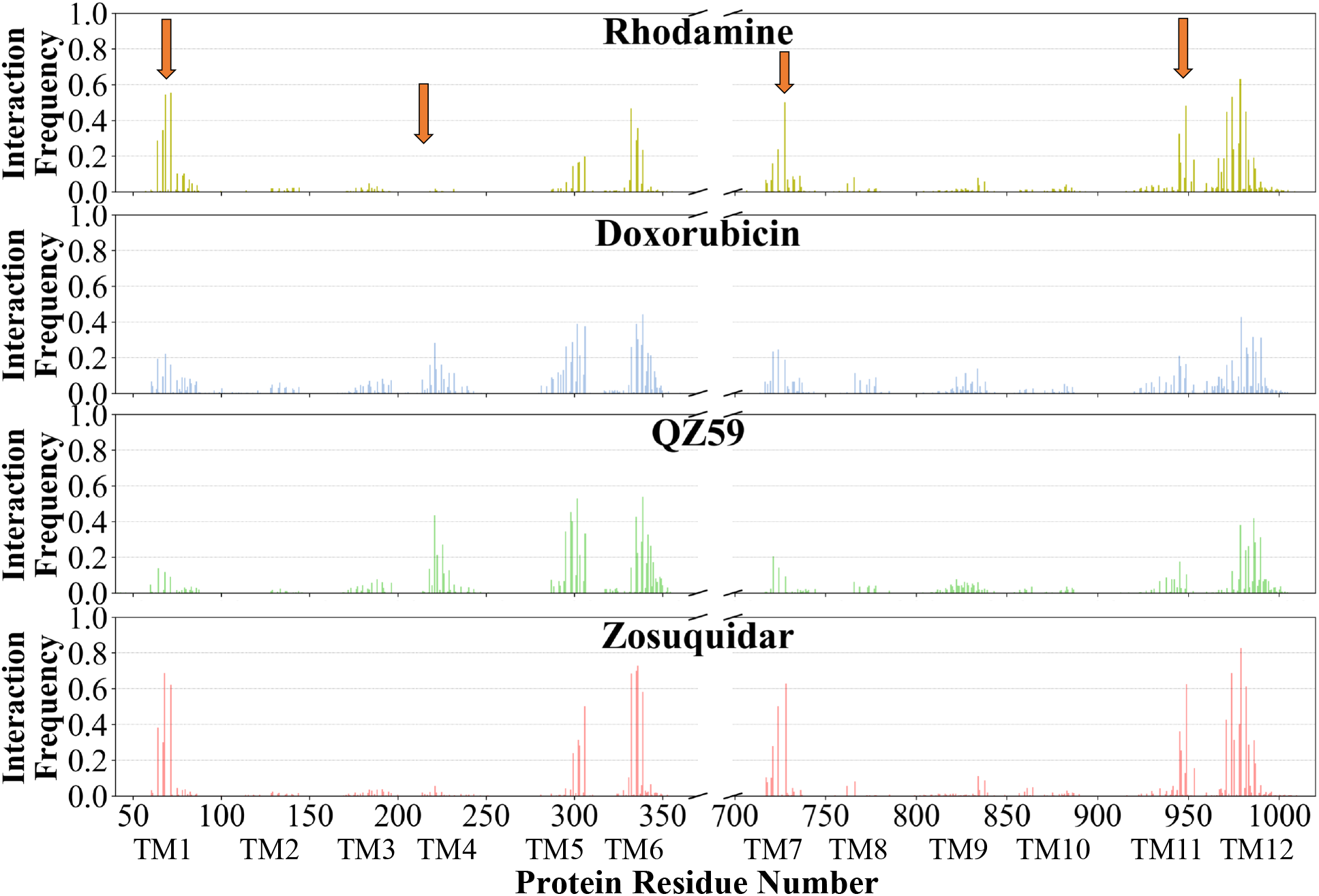
Interaction frequency of binding residues. The normalized interaction frequencies of the binding residues for all binding modes for 4 representative compounds are shown (data for the rest of the compounds are shown in Fig. S13). The residues are assumed to interact with the docked compound if their heavy atoms are within 4 Å. The orange arrows point to the regions of the protein showing differences in their interaction patterns for different compounds. High-affinity modulators like zosuquidar display the highest interaction frequencies with the binding residues pointing to a more specific mode of binding for these compounds.

Based on the most frequently (top 10) interacting residues, all compounds display binding preference for a set of core aromatic/hydrophobic residues present at the apex of the DBP: Y306, L335, I336, F339 and F979 (Fig. 9 and Table 2). High-affinity modulators and small substrates additionally show binding preference for a set of majorly aromatic residues, also lying close to the apex of the DBP: M68, F71, F332, F728, Y949, F974 and M982. Low-affinity modulator and large substrates on the other hand show preference for a different set of mostly hydrophobic residues of the DBP present in the TMD2: I221, I225, I295, I298, I299, M986 and M990. As high-affinity modulators/smaller substrates and low-affinity modulator/large substrates do not share any residue other than the core residues, these compounds likely bind to distinct but overlapping sites in Pgp.

**Figure 9:**
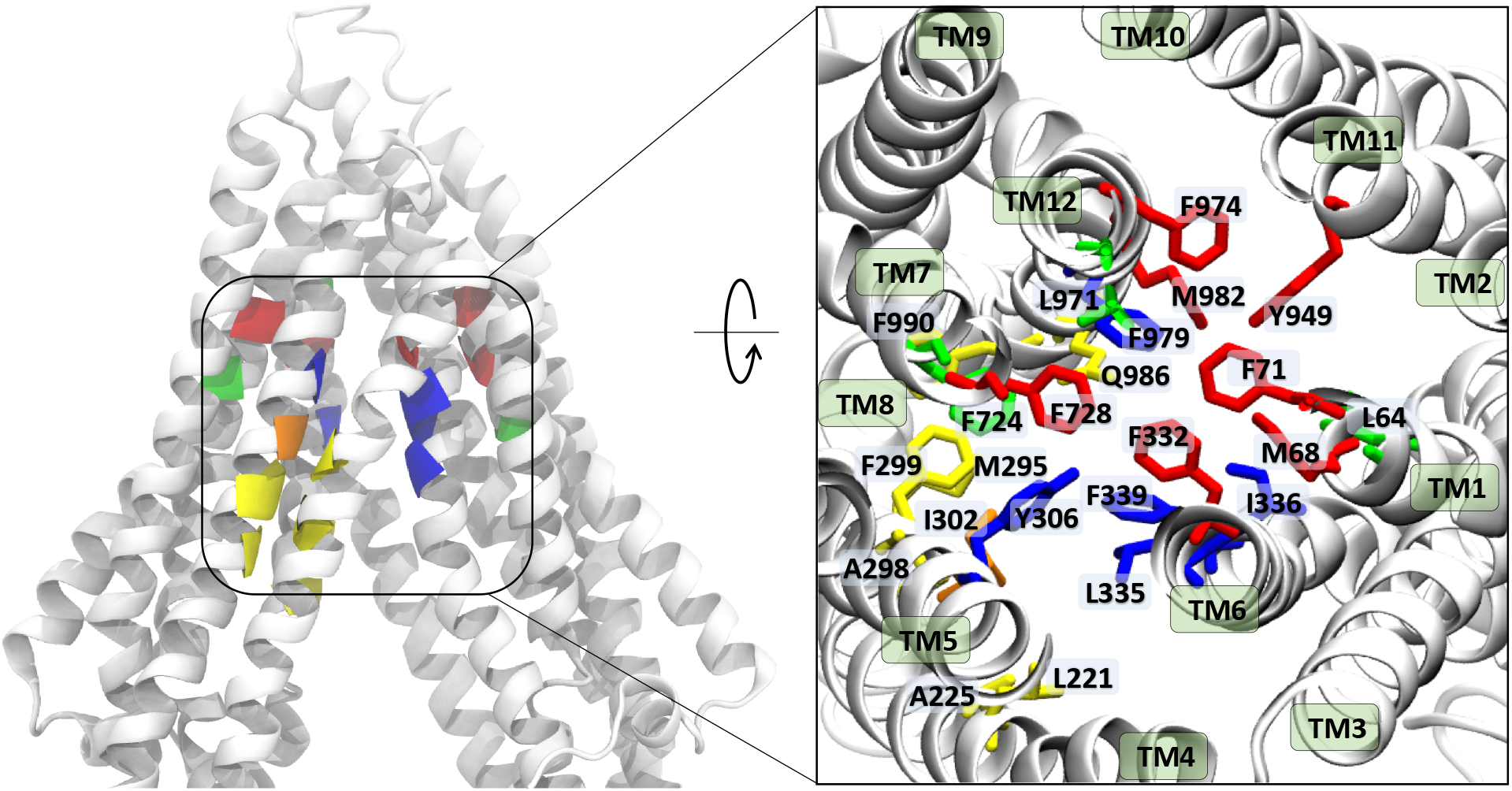
Ligand binding residues identified in Pgp. Binding site residues in Pgp showing the highest (top 10) interaction frequencies (from Fig. 8 and S13) with the tested compounds are shown in cartoon representation of the protein backbone (left) and in stick representation (inset, right). The different TM helices are labelled in the inset. The binding residues common to all compounds are shown in blue, residues common to binding of small substrates/high-affinity modulators are shown in red, residues common to low-affinity modulators/large substrates are shown in yellow, residues common to small substrates/low-affinity modulators are shown in orange, and residues showing preference for only small substrates are shown in green. High-affinity modulators share all binding site residues with small substrates and show binding preference for the M1 subsite, whereas low-affinity modulators and large substrates show binding preference for residues forming the M2 subsite, lying below the M1 subsite and partially overlapping with it.

**Table 2:** Top 10 binding site residues, based on the highest interaction frequencies, for each compound.

| **Compound** | **Top Binding Residues** |
| --- | --- |
| Colchicine | L64, M68, F71, F332, L335, I336, F728, F974, F979, M982 |
| Doxorubicin | I221, I299, I302, Y306, L335, I336, F339, F979, M986, M990 |
| Hoechst | F71, I302, Y306, F332, L335, F339, F728, Y949, F974, F979 |
| Prazosin | M68, Y306, F332, L335, I336, F339, F724, Y949, F974, F979 |
| Progesterone | M68, F71, Y306, F332, L335, I336, F339, F728, F974, F979 |
| Rhodamine | M68, F71, F332, I336, F728, Y949, F971, F974, F979, M982 |
| Verapamil | M68, F71, Y306, F332, L335, I336, F728, F974, F979, M982 |
| Vinblastine | I221, I225, I295, I298, I299, I302, Y306, F339, M986, M990 |
| QZ59 | I221, I295, I298, I299, I302, Y306, L335, F339, F979, M986 |
| Laniquidar | M68, F71, Y306, F332, L335, I336, F728, Y949, F974, F979 |
| Tariquidar | M68, Y306, F332, L335, I336, F339, F728, F974, F979, M982 |
| Zosuquidar | M68, F71, F332, L335, I336, F728, Y949, F974, F979, M982 |
| Diphenhydramine | M68, F71, F332, F724, F728, Y949, F953, L971, F974, F979 |
| Trimethoprim | F299, I302, Y303, Y306, F339, Q721, Q834, F979, A983, V987 |

### Differential binding of substrates, modulators and non-substrates

Ranking of the compounds based on their predicted binding affinities to the highly populated M1 site shows that the high-affinity modulators bind with relatively strongest affinities to the extended ensemble, whereas non-substrates display the weakest binding affinities (Fig. 10A). The substrates and low-affinity modulator lie between these two extremes. High-affinity modulators and most substrates also display high binding populations in the M1 site, ranging between 40-80% for the different compounds (Fig. 10B). In contrast, the low-affinity modulator and large substrates show low binding populations in the M1 site, and instead display higher binding populations in the second major binding site of the protein, the M2 site (Table 1).

**Figure 10:**
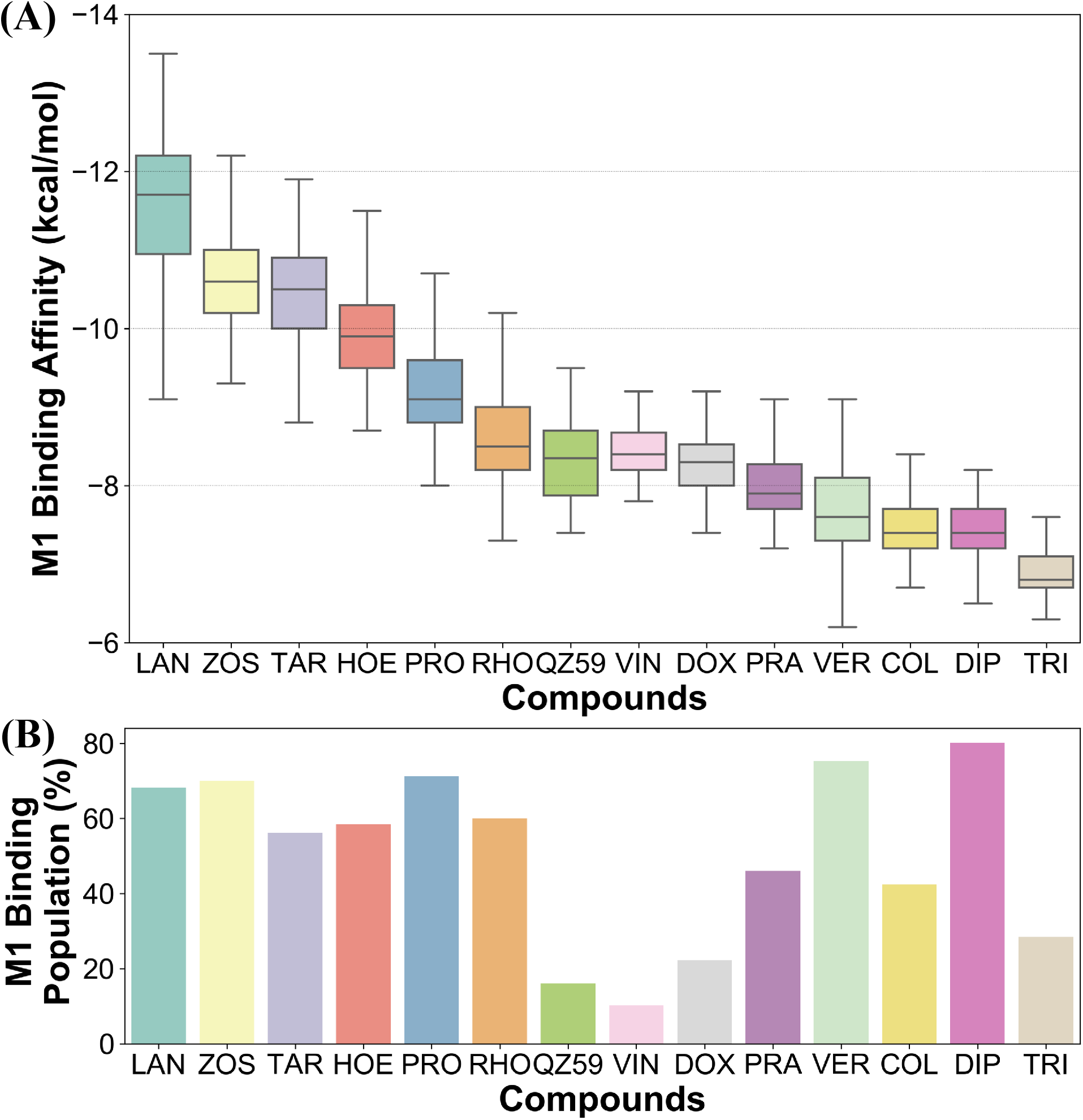
Differentiating between different classes of ligands. The binding site preference of different compounds was evaluated in terms of (A) the predicted binding affinities calculated for the extended ensemble of the protein in the M1 subsite, and (B) the respective binding populations in the same site. Ranking the compounds based on their binding affinities placed the high-affinity modulators and the non-substrates at the two extremes, with low-affinity modulators and substrates lying between them. Comparison of the binding site population in the M1 subsite further distinguished the high-affinity modulators and small substrates (showing high populations in M1 subsite) from low-affinity modulators and large substrates (showing higher populations in the M2 subsite instead).

## Discussion

We have put forth a protocol, termed as extended-ensemble docking, that can be used to characterize the binding of small molecules to proteins undergoing large-scale conformational changes (Fig. 2). Employing the technique to the case of ligand binding to multidrug resistance protein, Pgp, here we make use of a specifically designed SMD protocol utilizing system-specific CVs to describe the alternate-access mechanism of the protein and to generate an extended ensemble for it to be used in docking. Multiple runs with different orders of the CVs allowed us to identify a low-work transition pathway, corresponding to the most mechanistically probable transition pathway connecting the two major functional states of Pgp. The extended ensemble of the protein was then generated along this transition pathway followed by a series of docking and clustering calculations, allowing us to further characterize the different binding sites in Pgp as they evolve during the transition cycle of the transporter and the differential preference of compounds for these sites.

The global conformational changes during the transition cycle of Pgp are seen to modulate the size and shape as well as the ligand binding propensities of the binding pockets formed in the transporter (Fig. 4). Radioligand binding assays characterizing the binding of Pgp substrates as well as a study measuring drug binding affinity and reactivity to an antibody sensitive to Pgp conformational state, have shown that binding sites in Pgp display differential binding affinities for substrates depending on the conformation of the transporter during its catalytic cycle. ^75, 76^ In the present study, these differences are reflected in the large differences observed in the predicted binding affinities of compounds to the extended ensemble of the protein (Fig. 5). All compounds are seen to display stronger binding affinities to Pgp conformations other than its starting crystal structure, showing that experimentally determined protein structures may only reflect one of the many interaction possibilities and not necessarily the best one, between the transporter and its substrates.

Previous drug-drug interaction studies utilizing fluorescence spectroscopy,^77, 78^ kinetic studies of substrate binding and transport^79, 80^ as well as competitive binding assays^81, 82^ have postulated at least 3 binding sites in the TMDs of Pgp. These sites have been biochemically defined as H and R sites ^77, 78^ showing preference for substrates hoechst and rhodamine, respectively, and the modulatory M site showing preference for vinblastine^42, 81^ and verapamil, ^79, 80^ two compounds that have been described to function as both modulators and substrates of Pgp.^83–86^ A previous single-point docking study targeting the crystal structure of Pgp in the IF state assigned the above sites to different regions in the TMDs but showed that the docked compounds do not display binding preference for any specific site (i.e., the compounds showed similar binding affinities to all 3 sites). ^36^ In the present study, in contrast, we show that all compounds show binding preference for only one major substrate/modulator binding site, analogous to the M site, encompassing the top half of the DBP, whereas the sites present in the lower half of the DBP and in individual TMD leaflets, analogous to the H and R sites described above, show low binding populations and comparatively weaker binding affinities for all compounds. The H and R site may thus only function as poly-specific interaction sites in Pgp. We also report an interaction site on the membrane-facing surface of the TMD2, named as the S site, as well as additional sites formed on the extracellular side of TMDs forming as the transporter undergoes transition to the OF state, named as the E sites, that may assist in the extrusion of molecules out of the transporter.

Based on mapping of the clustering results onto the protein structure, the M site is found to consist of 3 subsites, which we name here as M1, M2 and M3. The M1 subsite is positioned in a central location at the apex of the TMDs, allowing interactions with residues from multiple helices from both TMD leaflets. The M1 subsite may represent the central binding site for substrate molecules (Fig. 6). High-affinity modulators are also predicted to bind to M1 but with comparatively stronger binding affinities and higher interaction frequencies than substrates, pointing to a more selective binding mode for these compounds compared to the substrates (Fig. 7 and Fig. 8). These modulators, showing IC_50_ values in the nanomolar range, ^87–89^ can also function as strong third-generation inhibitors of Pgp and may compete with low-affinity substrates binding to this site. Substrates verapamil and progesterone are seen to bind to the M1 subsite with high interaction frequencies comparable to high-affinity modulators, but with relatively weaker binding affinities. These molecules show IC_50_ values in the micromolar range and were previously characterized as first-generation inhibitors of Pgp.^90^ They require high concentrations to fully inhibit the transporter, to the degree that they start to display toxicity and thus not suitable for administration. ^91^ A recently published cryo-EM structure shows binding of a substrate molecule taxol to the M1 subsite, ^92^ whereas another structure shows two zosuquidar molecules bound to the M site of Pgp, with the first molecule occupying the M1 subsite and the second occupying the M2/M3 subsites. ^57^ These structural studies provide further evidence that the M1 subsite described in our study may represent the central substrate/modulator binding site in Pgp.

Larger low-affinity modulator QZ59 shows preference for the M2 subsite, with some binding populations also observed in the M1 and M3 subsites. Previously published crystal structure of Pgp with this modulator shows two copies of it bound to Pgp, with the first copy located in a site similar to the M2 subsite described in our study (partially within the TMD2 leaflet of Pgp), and the second copy located in the M1 subsite. ^28^ In multiple crystal structures of Pgp in complex with different structural analogs of QZ59, simultaneous binding of two ligands have also been observed to the M2/M1 or M3/M1 subsites of Pgp.^51^ Larger substrates like doxorubicin and vinblastine also show a similar binding behavior to the low-affinity modulator, with a preferential binding for the M2 subsite though some binding populations were also observed for the M1 and M3 subsites. Similar to the QZ59 analogs, it is possible that two (or more) of these substrates may more stably bind together to the multiple M subsites.

Mapping of the clustering results on the protein structure further showed that the H and R sites identified in the separate TMD leaflets (showing much lower binding populations compared to the M site), each consisted of two subsites, H1 and H2, and R1 and R2, respectively (Fig. 6). The H1 and R1 subsites are found in open IF-like conformations and may be involved in formation of initial interactions with the molecules as they partition from the membrane through the two proposed drug-entry portals located on either sides of Pgp.^49^ Increase in the local concentration of molecules at these interaction sites may facilitate their eventual movement towards the central binding site (M1). The subsidiary S site (consisting of the S1 subsite) on the membrane facing side of TMD2 may allow formation of encounter complexes of substrates with the transporter, and it may function as an entry site for these molecules into Pgp. A similar site has also been identified in the crystal structure of Pgp in complex with QZ-Valine, ^51^ a derivative of QZ59, as well as in MsbA (a bacterial homolog of Pgp) by using site-specific spin labels. ^93^

The extracellular E site, consisting of E1 and E2 subsites, represents novel binding sites identified in this study that partially overlap with the M1 subsite in the DBP and are formed as the TMDs open to the extracellular side during the formation of the OF state. These sites may play a role in assisting the diffusion of the substrate out of the transporter, as the DBP is exposed to the extracellular environment in the OF state, by providing additional interaction sites.

Experiments using photoaffinity labelling of Pgp substrates as well as crystal structure studies have suggested the presence of high- and low-affinity binding sites in Pgp.^51, 94, 95^ Potential of mean force calculations using umbrella sampling also identified multiple overlapping binding locations for different substrates inside the TMDs of Pgp.^96, 97^ The results of our extended-ensemble docking provide further evidence for this proposal, where the M1 and M2 subsites serve as binding sites for the high-affinity modulators/small substrates and large, low-affinity modulators/substrates, respectively (Fig. 11A). These subsites form overlapping sites within the DBP of Pgp, with all modulators and substrates showing preference for the same set of core residues (Fig. 9). The low-affinity binding sites proposed in the above studies may comprise of the H, R and S sites located below the M site in the individual TMDs of Pgp. Correspondingly, the predicted binding affinities display an increasing trend between these low- and high-affinity binding sites (Fig. 11B), likely promoting the diffusion of molecules towards the central binding site (M1) and their eventual extrusion through the extracellular sites.

**Figure 11:**
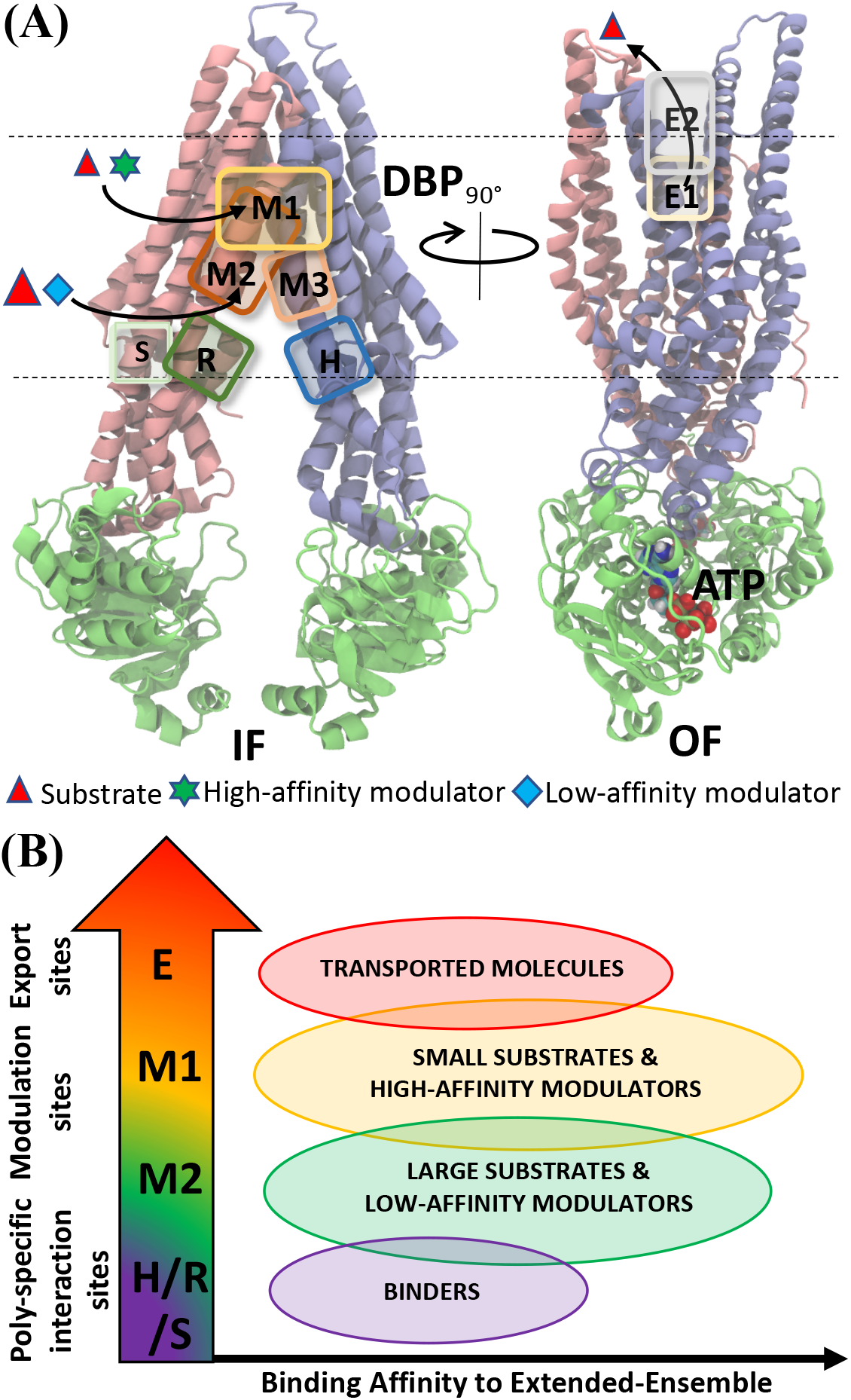
Ligand binding sites in Pgp identified by extended ensemble docking. (A) Different binding sites are shown overlaid with representative IF (left) and OF (right) conformations. The two transmembrane leaflets are shown in blue (TMD1) and pink (TMD2) cartoon representation, respectively, and the NBDs are shown in green. The major substrate-modulator binding site (M1), as well as the low-affinity modulator/large substrate binding site (M2) are observed throughout the conformational transition of Pgp, whereas extracellular sites (E1 and E2) are only observed in the OF-like states. (B) By combining the predicted binding affinities of all compounds in the different binding sites obtained in the extended ensemble (Fig. 7 and S12), we observe differences in the relative binding affinities of the poly-specific interaction sites (H/R/S), modulation sites (M1/M2) and the extracellular sites (E). These differences may facilitate the transport of the molecules from inside to the outside of the cell.

The differential binding behavior of the compounds to the above subsites in the extended ensemble of the protein further allowed us to differentiate between different classes of compounds (Fig. 10). The availability of different binding sites, modulated by the conformational changes in the transporter, as well as modulation of their binding propensities may form the basis of poly-specificity of Pgp towards different substrates. Furthermore, information on molecules showing stronger binding affinity (specificity) to the protein extended ensemble instead of a single structure, may provide a better framework for the development of specific, fourth-generation inhibitors of Pgp.

## Conclusion

Large flexible protein like Pgp have proven to be difficult drug-targets; a number of recent ventures in developing new drug molecules targeting Pgp have failed. Further optimization strategies using crystal or cryo-EM structures can be rather difficult, as these proteins constantly fluctuate between different conformational states, concurrently leading to changes in the available/characterized binding sites. This challenge also represents a major roadblock for developing accurate structural-activity relationship models for conformationally heterogeneous proteins. Future endeavours in structure-based drug discovery targeting these proteins mandates new in-silico strategies, taking into account the inherent flexibility and inter-talk between the different domains of the protein as it undergoes transition between its major functional states. We have described here a new docking protocol pertaining to these systems, allowing high-throughput virtual screening of an extended ensemble derived along the protein’s transition pathway, instead of targeting a single crystal/cryo-EM structure. This has allowed us to provide novel insights into the differential binding behaviors of different classes of compounds in Pgp. Future applications to other flexible proteins systems may open a new frontier in in-silico drug design targeting these important drug targets.

## Supporting information

Supplementary figures and results

## Acknowledgement

The authors acknowledge support by the National Institutes of Health under award numbers P41-GM104601 and R01-GM123455 (to E.T.). We also acknowledge computing resources provided by Blue Waters at National Center for Supercomputing Applications (NCSA).

