## Supplementary figures and results for "Extended-Ensemble Docking to Probe Evolution of Ligand Binding Sites During Large-Scale Structural Changes of Proteins"

### Supplementary Information

Here we provide more details on the SMD results used for generating the extended ensemble of Pgp.

#### Conformational ensemble of Pgp during IF to OF transition

Starting from 20 pre-equilibrated IF structures of Pgp and using the modeled OF structure as the target, SMD simulations were performed using different transition protocols representing mechanistically distinct transition pathways. The RMSD of the starting equilibrated IF structures highlighted the conformational heterogeneity of the IF state of the transporter (Fig. S5). The comparison of the non-equilibrium work profiles shows a consistent trend emerging from the 4 transition protocols applied. The SMD simulations utilizing protocol P4 ($\alpha$ + NBDi + SB + $\beta$) show an overall lower non-equilibrium work values compared to the other 3 protocols, for the 20 independent runs conducted for each protocol (Fig. S4A). Furthermore, final structures generated using this protocol show lower RMSD values with respect to the target OF structure compared to other protocols (Fig. S4B). The transition pathway obtained from protocol P4 was thus considered to be the most probable mechanistic pathway connecting the two functional states of Pgp and was further utilized for longer SMD runs.

The longer SMD simulation showed lower overall work required for the transition compared to the shorter SMD runs (Fig. S4A and Fig. S6A). This is expected as longer timescales allow the system to remain closer to its ideal pathway, decreasing dissipation and leading to lower work values. Concurrently, a more linear change in the CVs over time is observed (Fig. S6B and C). The different domains (TMDs, NBD1, NBD2) of the protein generated at the end of the SMD simulation showed RMSD values ranging between 3 and 5 Å with respect to the target OF structure (Fig. S7A) and between 2 and 6 Å for the individual TM helices (Fig. S7B), reiterating the highly flexible nature of Pgp as well as possible differences between the OF state and the homologous structure used for the construction of the OF model. The ensemble of Pgp structures for subsequent docking calculations was obtained along the transition path by selecting 50 snapshots at regular time intervals spanning nearly equally the conformational space sampled by the TMD intracellular and extracellular angles and the distance between the two NBDs (Fig. S8).

Table S1: The distance CV parameters used for applying salt-bridge interaction (SB) and NBD interactions (NBDi; comprising ATP and X-loop interactions) during IF to OF SMD simulations.

| | **Atom-1** | | **Atom-2** | | **Target** |
|---|---|---|---|---|---|
| **CV** | **Residue** | **Atom** | **Residue** | **Atom** | **Distance (Å)** |
| SB | K185 | NZ | D993 | OD1/OD2 | 2.8 |
| NBDi | G530/G1175 | N | ATP | O1G | 2.76 |
| NBDi | S528/S1173 | OG | ATP | O1G | 2.88 |
| NBDi | S528/S1173 | OG | ATP | O3B | 3.30 |
| NBDi | Q531/Q1176 | OE1 | ATP | O2' | 2.80 |
| NBDi | L527/L1172 | CA | ATP | C8 | 4.21 |
| NBDi | D163 | OD1/OD2 | K1167 | NZ | 2.8 |
| NBDi | E522 | OE1/OE2 | T806 | HG1 | 2.8 |

Table S2: Binding pocket size and propensity of ligand binding (PLB) to the extended-ensemble of Pgp.

| | **TMD-Apex** | | **TMD1** | | **TMD2** | | **TMD-Ex.*** | | **TMD-In.*** | |
|---|---|---|---|---|---|---|---|---|---|---|
| **Structure** | **Size** | **PLB** | **Size** | **PLB** | **Size** | **PLB** | **Size** | **PLB** | **Size** | **PLB** |
| Crystal structure | 219 | 5.4 | 47 | 0.4 | 84 | 0.6 | - | - | - | - |
| Snapshot-1 | 187 | 4.7 | 89 | 1.3 | 54 | 1.3 | - | - | - | - |
| Snapshot-10 | 210 | 3.7 | 76 | 1.1 | 117 | 1.4 | - | - | - | - |
| Snapshot-20 | 327 | 7.1 | 53 | 0.6 | 154 | 1.9 | - | - | - | - |
| Snapshot-30 | 473 | 7.8 | 111 | 0.5 | 67 | 0.3 | - | - | - | - |
| Snapshot-40 | 232 | 4.4 | 94 | 1.4 | 75 | 0.6 | 77 | 0.6 | 144 | 0.6 |
| Snapshot-50 | 249 | 0.7 | - | - | - | - | 49 | 0.05 | - | - |

* TMD-Ex. and TMD-In. represent the binding pockets predicted on the extracellular and intracellular sides of TMD.

Table S3: Percentage distribution of binding modes of all compounds in different binding clusters and their correspondingly mapped sites in the protein.

| **Cluster Number** | **Mapped Protein site** | **Percentage** |
|---|---|---|
| 1 | E1 | 0.9 |
| 2 | E3 | 0.8 |
| 3 | E2 | 6.1 |
| 4 | M1 | 50.4 |
| 5 | M2 | 27.8 |
| 6 | R2 | 2.4 |
| 7 | R1 | 3.3 |
| 8 | S1 | 2 |
| 9 | S2 | 0.7 |
| 10 | H1 | 2.7 |
| 11 | M3 | 2.2 |
| 12 | H2 | 0.7 |

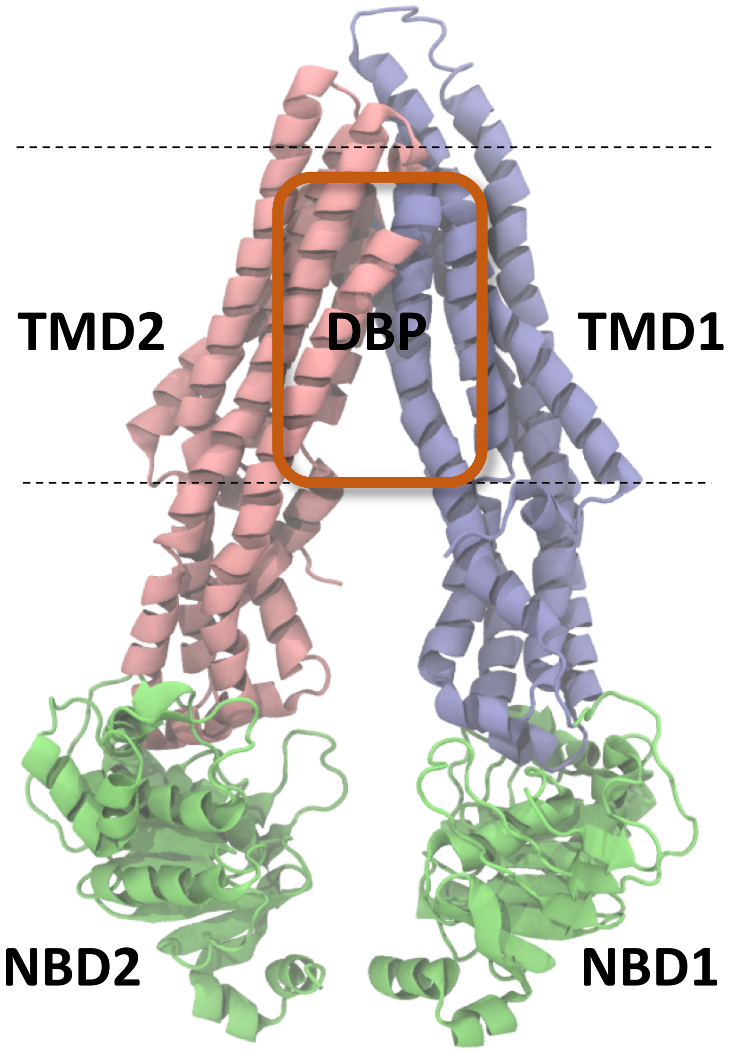


Figure S1: **Structure of Pgp.** Pgp is shown as cartoon representation. The multi-domain protein consists of two transmembrane (TMD) domains, TMD1 and TMD2, shown in blue and pink, respectively, each connected to a nucleotide binding domain, NBD1 and NBD2, respectively, shown in green. A large drug binding pocket (DBP) in the TMD allow binding of different classes of molecules.

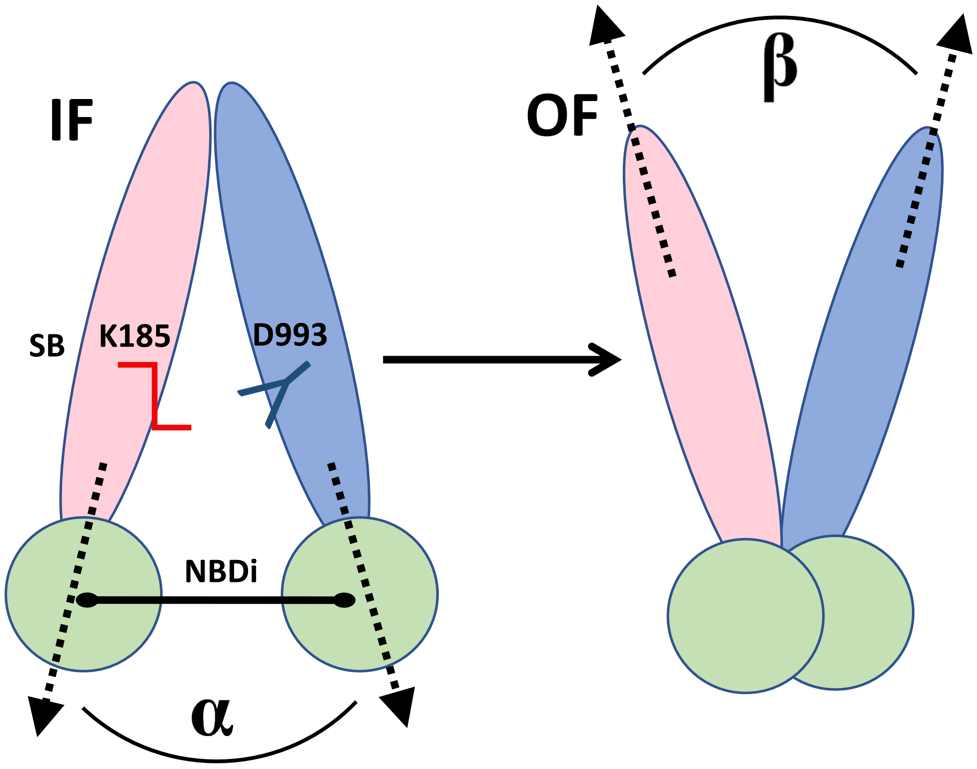


Figure S2: **CVs used to induce structural transition of Pgp between conformational state IF and OF.** The two TMD leaflets of Pgp, each connected to an NBD, are shown in pink and blue, respectively. The four system-specific CVs, $\alpha$, (opening/closing of the cytoplasmic side) $\beta$ (opening/closing of the extracellula side), NBDi (contacts between the NBDs), and SB (salt bridge between K185 and D9993), used to steer the conformational transition of the protein, following the 'alternate-access' mechanism of ABC-transporters, are shown. $\alpha$ and $\beta$ describe the intracellular and extracellular gates in the TMDs, respectively, NBDi describes the ATP and X-loop interactions in the NBDs necessary for the formation of the NBD dimer, and SB describes a salt-bridge formed within the TMD.

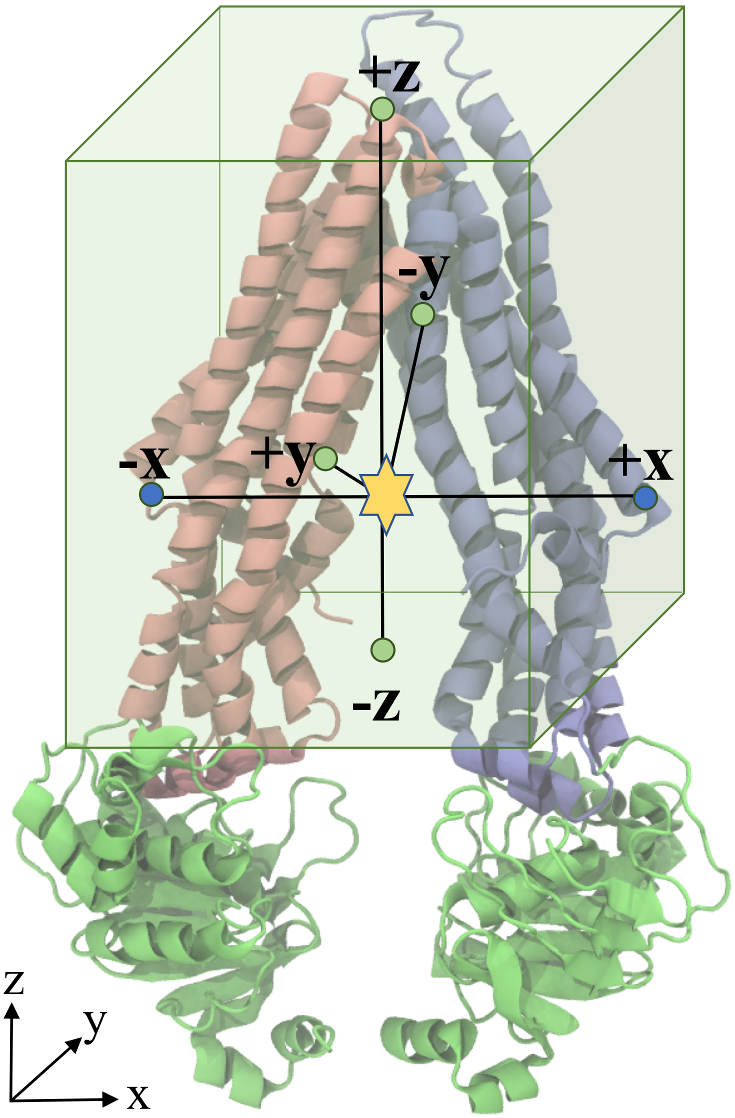


Figure S3: **Distance used for clustering.** Distances describing each binding mode (yellow star) to the TMDs were calculated with respect to 6 points in the space. Four fixed points in the $+y$, -$y$, $+z$, and -$z$ directions (shown as green dots) were selected as the centers of the four side faces of the docking grid-box around the TMD (green transparent rectangular prism), respectively, and two variable points in the $+x$ and -$x$ directions (shown as blue dots) were selected as the C-$\alpha$ atoms of first residues of TM1 and TM7 helices, respectively.

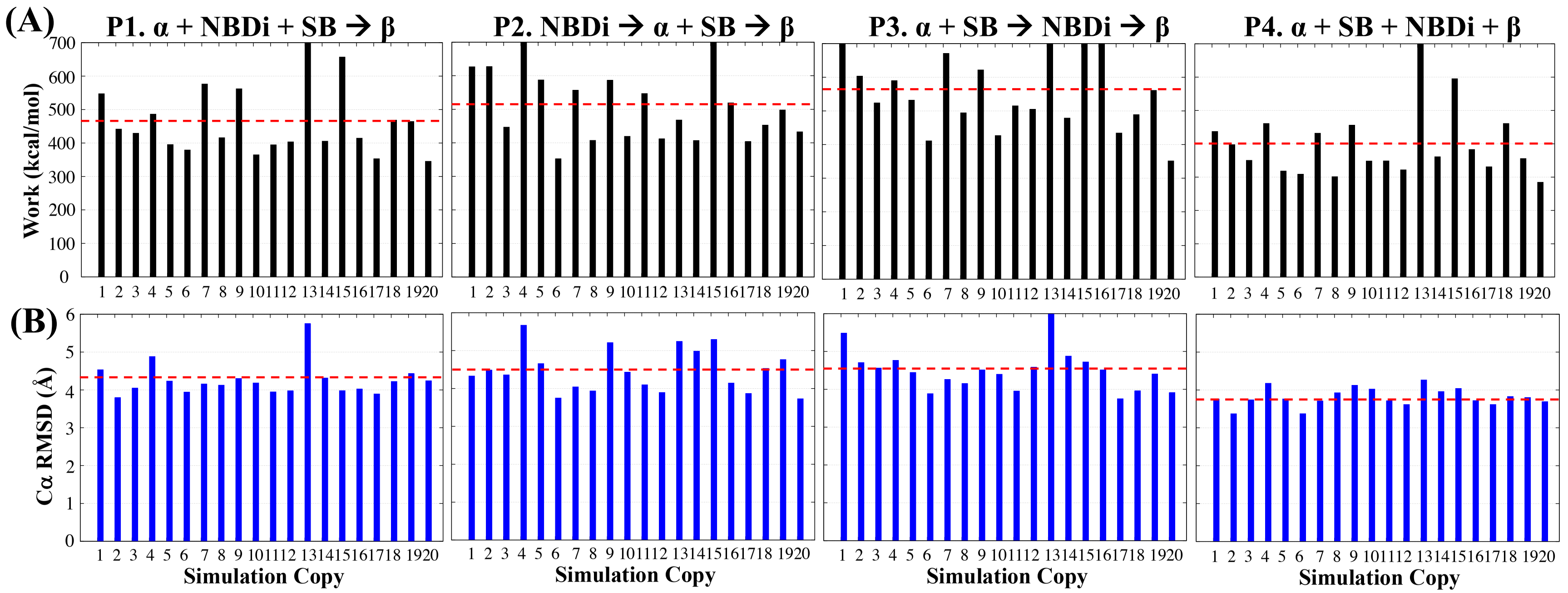


Figure S4: **Non-equilibrium work relations and structural variation with respect to target structure in SMD simulation.** A) The non-equilibrium work profiles for the 4 transition protocols (P1, P2, P3 and P4), each defined by a different order of applied four CVs, used in SMD simulations for transition between the IF and OF states of Pgp are provided. The work values were calculated for 20 independent 30-ns SMD runs for each protocol, each starting from a separate IF structure selected from an initiail equilibrium pool. B) The Cα RMSD with respect to the target OF structure for the above SMD runs are shown. The dashed red lines in (A) and (B) represent the average non-equilibrium work or RMSD, respectively, calculated for the 20 independent runs in each protocol. Overall, P4 showed the lowest work values for the transition, with the final structure reaching closest (in terms of RMSD) to the target OF structure.

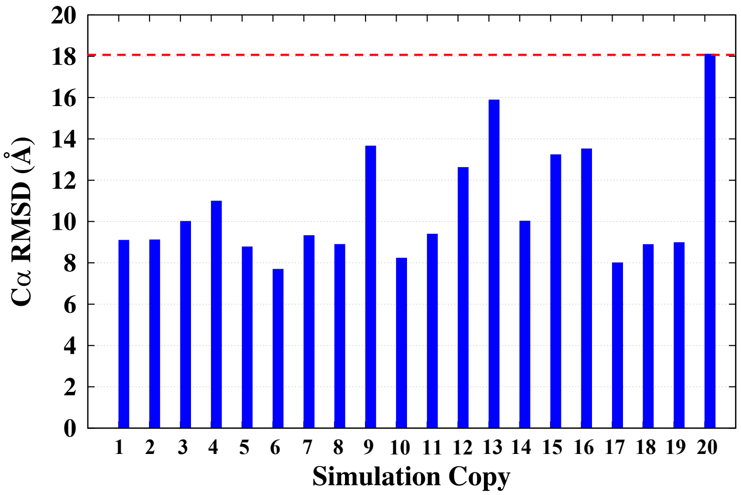


Figure S5: **Conformational heterogeneity of the IF state.** The plot shows C$\alpha$ RMSD for each of the 20 starting IF structures with respect to the target OF structure. The dashed red line represents the largest RMSD, corresponding to the equilibrated IF structure 20 which was selected as the starting structure for the longer SMD run used for the generation of the extended ensemble.

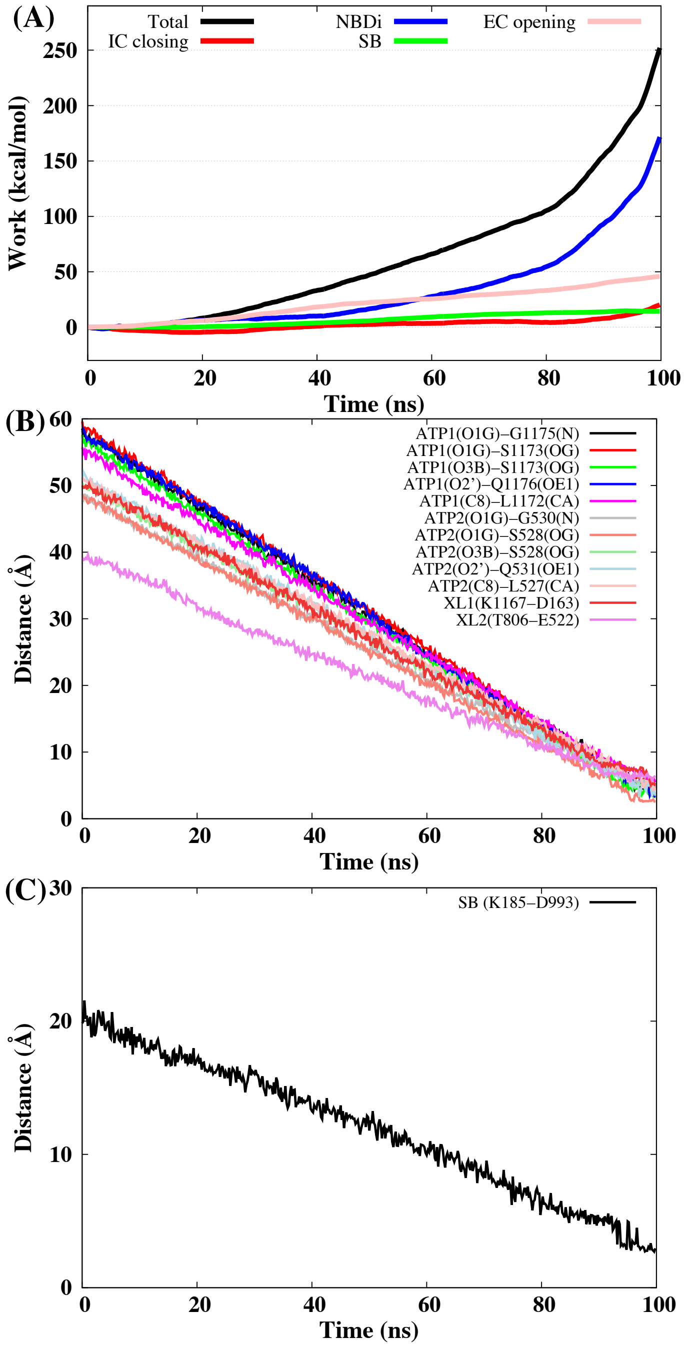


Figure S6: **CVs during SMD simulations.** A) The total non-equilibrium work for the 100-ns SMD run is shown for the most efficient transition protocol (P4), along the work contributions along individual CVs. The longer SMD simulation showed lower overall work values compared to the shorter runs employing the same starting IF structure (Fig. S4). The changes in the values of the CVs comprising B) ATP-NBD and X-loop interactions (NBDi), and C) salt-bridge formation (SB) over the 100-ns SMD simulations are provided. Both these CVs show a linear decrease over the course of the simulation.

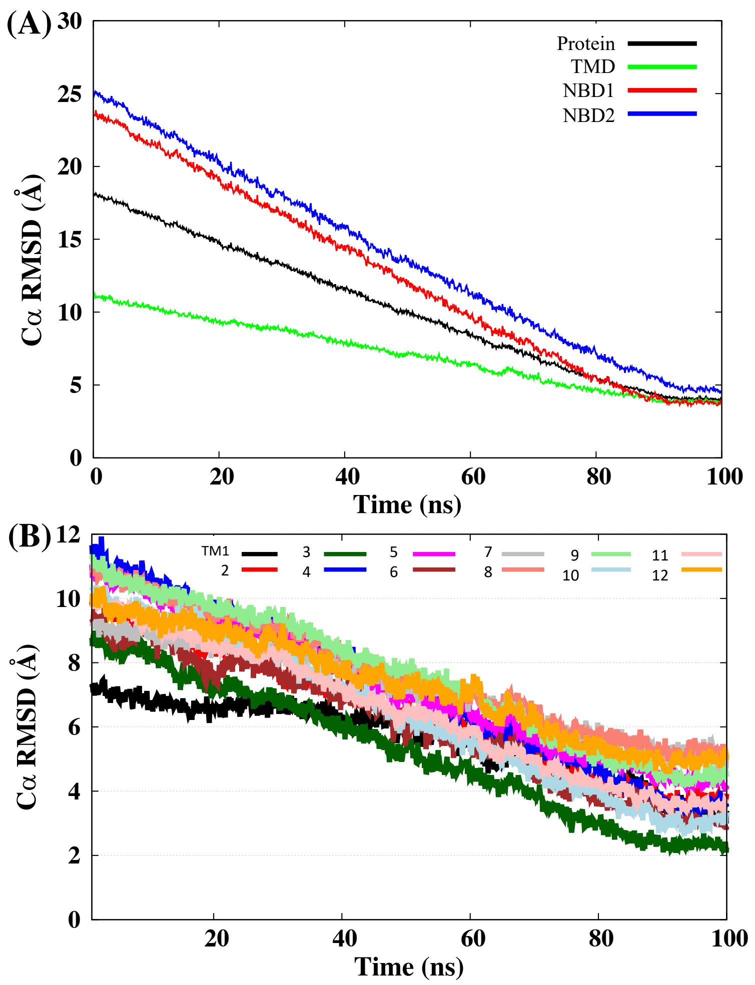


Figure S7: **Structural changes during SMD simulations.** A) The RMSD values for the whole protein and for the separate domains of Pgp are shown. RMSDs for TMD (TMD1+TMD2), NBD1 and NBD2 were calculated by first superimposing the trajectory using the whole protein backbone and then calculating the RMSD for the individual domains. B) The RMSD values of the individual TMD helices (TM1-TM12) with respect to the final OF structure are shown. These values were calculated by first superimposing the trajectory using the TMD of the protein and then calculating the RMSD for the individual TMs. Overall, the IF structure reached within ∼4 Å of the target OF structure.

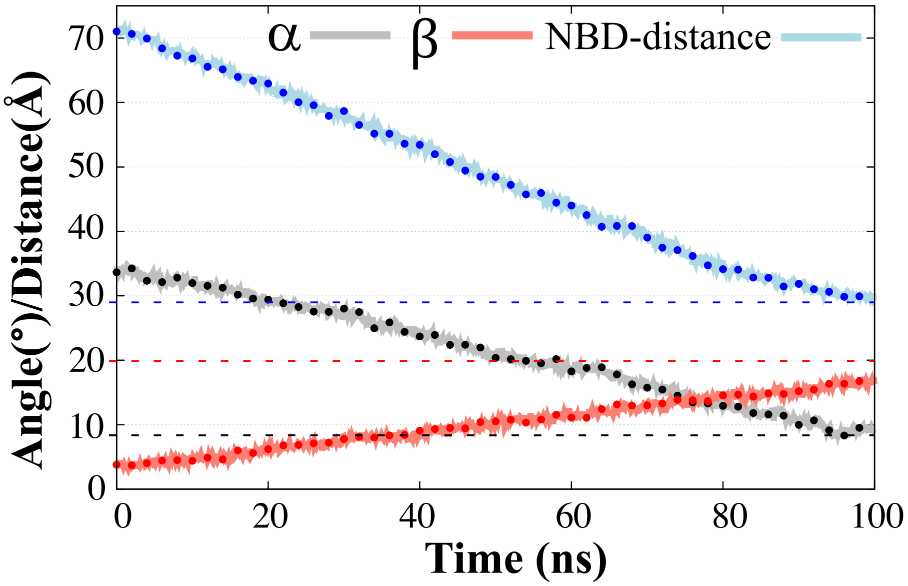


Figure S8: **Conformation selection of Pgp's extended ensemble.** A) The global conformational changes in Pgp during IF to OF transition were characterized in terms of intrcellular closing angle ($\alpha$), extracellular opening angle ($\beta$) and the distance between the centers of masses of the two NBDs. The extended-ensemble of the protein was generated by taking 50 snapshots equally distributed along this conformational space (shown as dots). The $\alpha/\beta$ angles and NBD-distance values in the target OF structure are shown as dashed lines of similar colors.

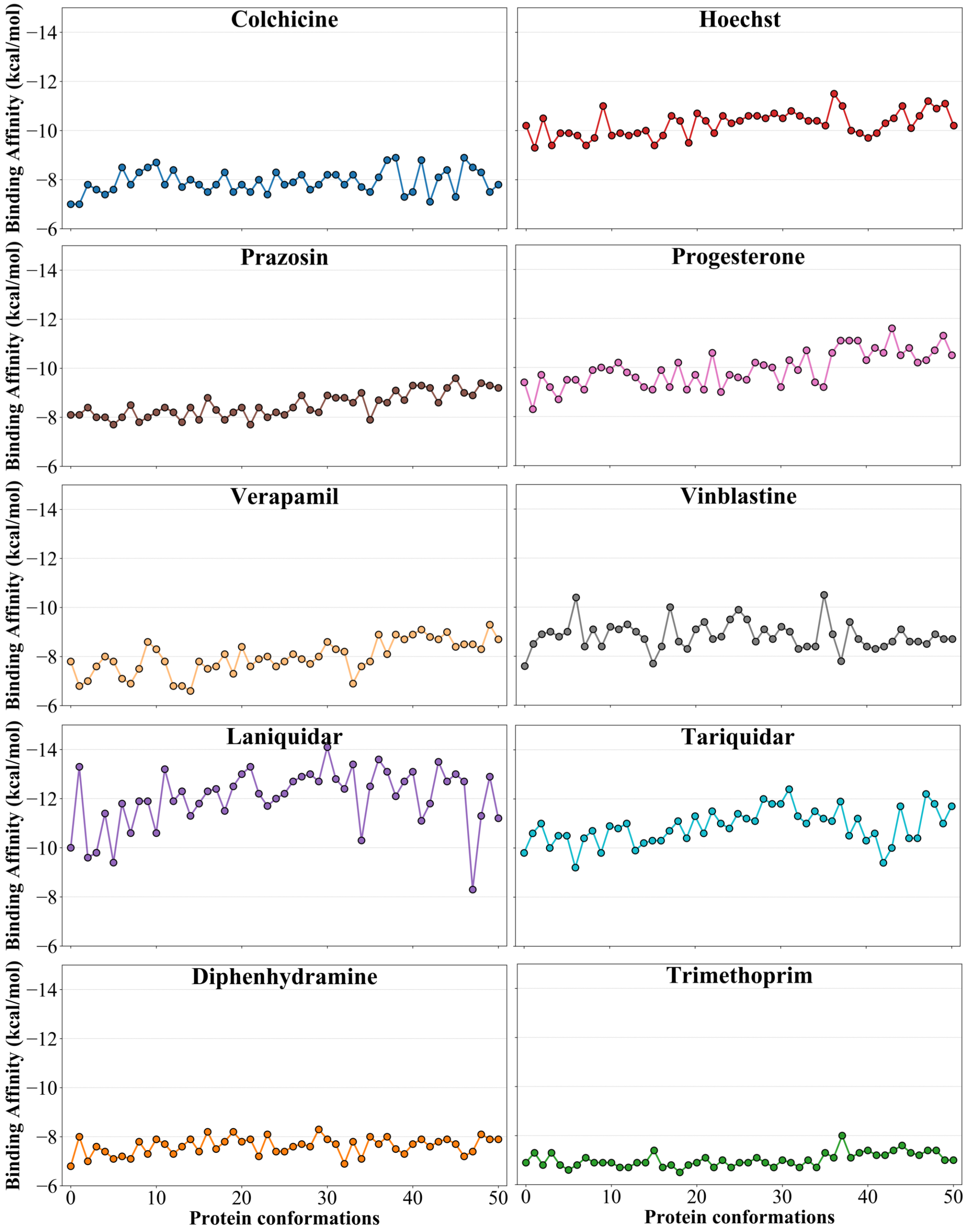


Figure S9: **Binding affinities to the extended-ensemble.** The predicted best binding affinities for all compounds to each of the selected 51 Pgp conformation (0-50) are shown. The $0^{th}$ conformation represents the starting crystal structure in the IF state used in generating the transition pathway.

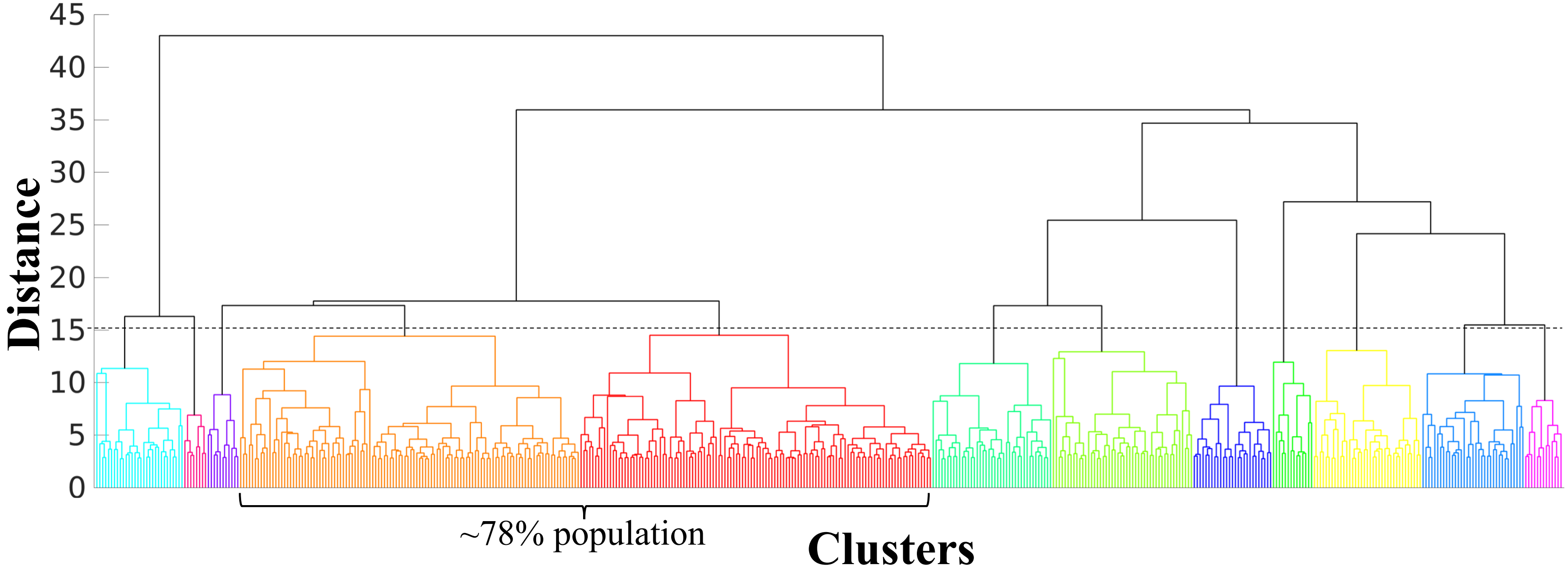


Figure S10: **Clustering Dendrogram.** Hierarchical tree or dendrogram generated using average linkage clustering method for clustering the distances of the binding modes for all docked compounds is shown. The cutoff distance (15) selected for clustering the dendrogram is shown as dashed black line. All tree nodes lying below the cutoff distance represent a separate cluster shown in a different color. The respective binding populations of the clusters are provided in Table S3, and the mapped binding sites on Pgp structure represented by these clusters are shown in Fig. 6 and Fig. S11. Clusters 4 and 5 together constituted 78% of all predicted binding modes.

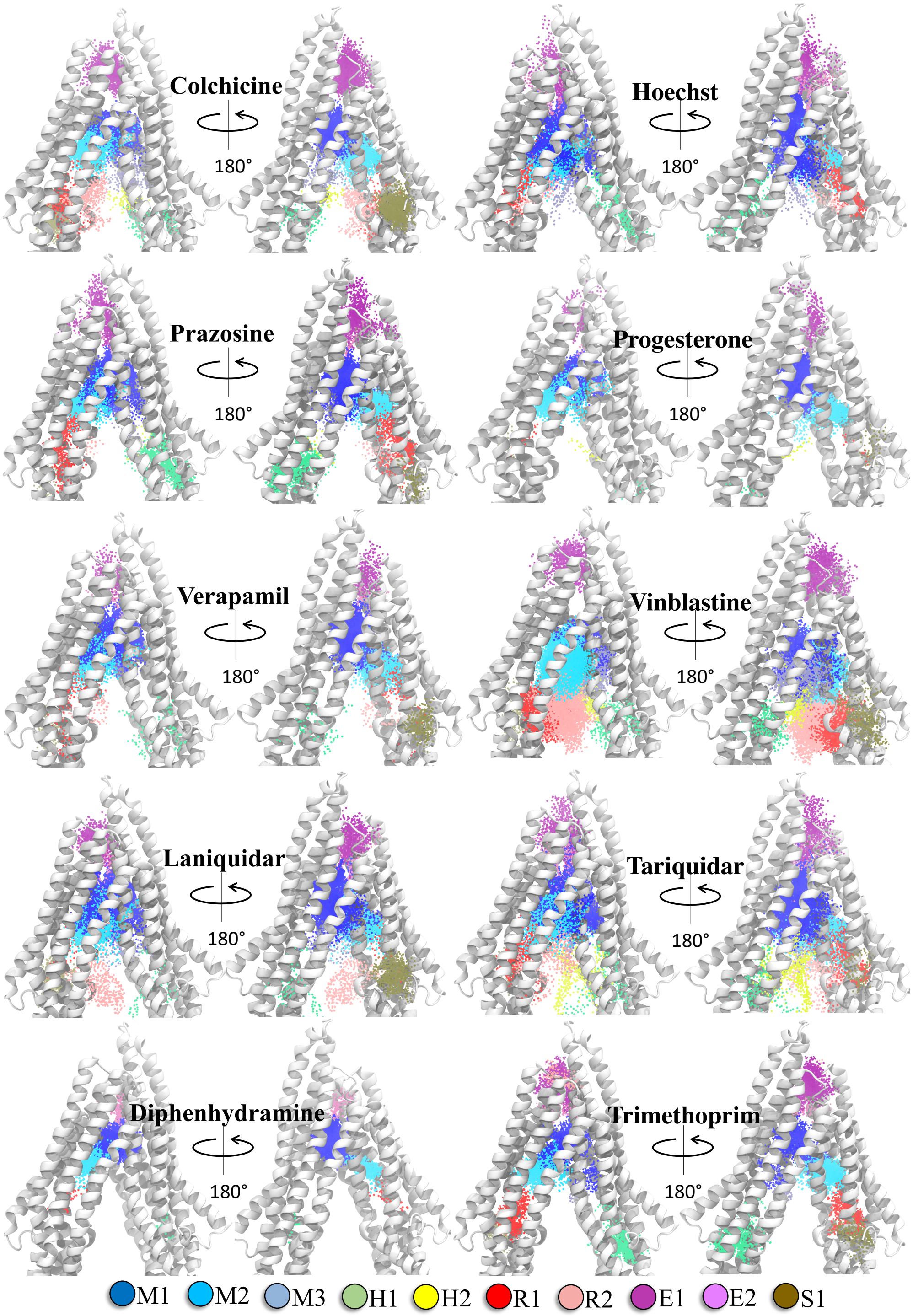


Figure S11: **Clustering of binding modes generated from extended-ensemble docking.** Clustering of the binding modes for all compounds in the extended ensemble of Pgp are overlaid onto a representative (IF) conformation of Pgp. Each binding cluster is shown in a different color with heavy atoms of the bound ligand shown as points. The density of points in each cluster represents the cluster population. The corresponding binding subsites in the protein are indicated in the legend at the bottom.

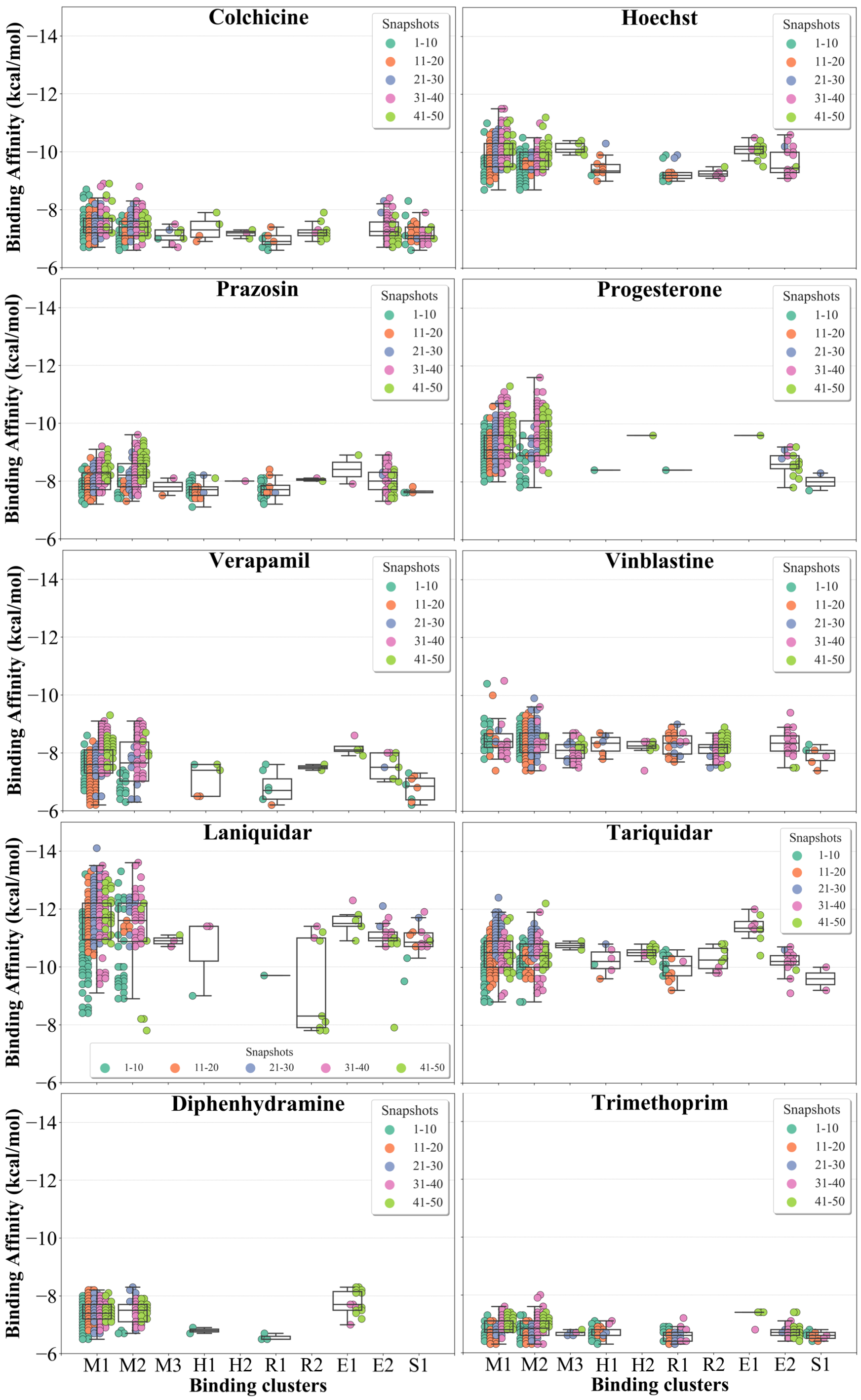


Figure S12: **Cluster binding energies.** The binding affinities of members (binding modes) of each cluster are shown as a swarm plot for different compounds. The snapshots belonging to different protein conformations are shown in different colors (defined in legend). Additionally, a boxplot, providing the median cluster values, Q1 and Q3 quartiles, as well as minimum (Q1 − 1.5 × interquartile range) and maximum (Q3 + 1.5 × interquartile range) binding affinity values, is overlaid on top of the swarm plot for each cluster.

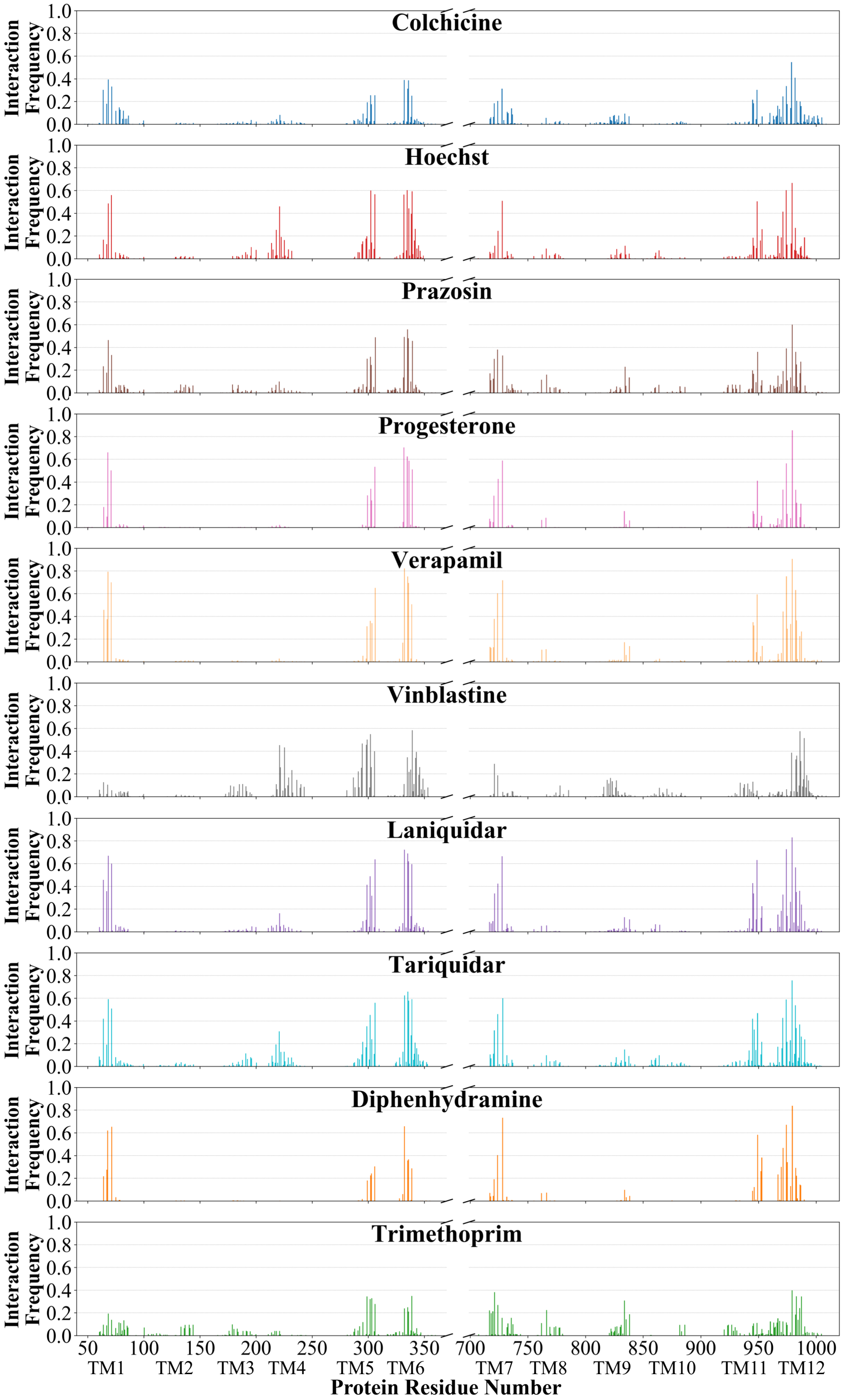


Figure S13: **Interaction frequency of binding residues.** The normalized interaction frequencies of the binding residues for all binding modes of different compounds docked to the extended-ensemble of Pgp are shown. The residues are assumed to interact with the docked compound if thier heavy atoms are within 4 Å.
